# Striatal output regulates the postnatal maturation of cortical circuits

**DOI:** 10.1101/2024.05.10.593512

**Authors:** Michael Janeček, Tara Deemyad, Yi-Chun Shih, Vicente Valle, Andrew D’Agostino, Michael Matarazzo, Megan S. Perez, Kyle D. Ketchesin, Susana da Silva, Rui T. Peixoto

**Affiliations:** Department of Psychiatry, University of Pittsburgh, 450 Technology Dr, Pittsburgh, PA 15219, USA; Center for Neuroscience at the University of Pittsburgh; Department of Ophthalmology, University of Pittsburgh, 1622 Locust St, Pittsburgh, PA 15219, USA; Department of Human Genetics, School of Public Health, University of Pittsburgh, Pittsburgh, USA; Department of Bioengineering, Swanson School of Engineering, University of Pittsburgh, Pittsburgh, USA; Department of Otolaryngology, Johns Hopkins School of Medicine, 720 Rutland Ave, Baltimore, MD 21205, USA

## Abstract

The dorsomedial prefrontal cortex (dmPFC) is interconnected with the basal ganglia (BG) through large-scale circuit loops that regulate critical motor and cognitive functions. In mice, these circuits undergo extensive postnatal maturation with marked changes in neural activity and expansion of synaptic connectivity. While cortical activity is known to regulate the development of downstream striatal circuits, the role of the basal ganglia in cortical maturation remains unknown. Here, we used mesoscale two-photon microscopy and whole-cell electrophysiology to examine whether striatal output during early postnatal development impacts the activity and maturation of upstream dmPFC circuits. We found that ablating spiny projection neurons of the direct or indirect pathways of the striatum during the first two postnatal weeks causes bidirectional changes in dmPFC neural activity, similar to observations in mature circuits. In addition, these manipulations alter the maturation of synaptic connectivity of dmPFC layer 2/3 pyramidal neurons, shifting the balance of excitation and inhibition of cortical circuits. These findings demonstrate that striatal output modulates the activity of cortical circuits during early postnatal development and suggest a regulatory role of the basal ganglia in the establishment of cortical circuits.

## Introduction

The dorsomedial prefrontal cortex (dmPFC) and the basal ganglia (BG) are functionally interconnected through large-scale cortico-BG-thalamocortical (CBGT) circuits crucial for motor control and cognition^1^. The striatum, the primary input nucleus of the BG, receives convergent excitatory input from cortex and thalamus, which synapse onto two classes of spiny projection neurons (SPN) that establish functionally antagonistic outputs termed direct (dSPN) and indirect (iSPN) pathways. In the mature brain, activation of dSPNs increases cortical activity, while activation of iSPNs suppresses it^2–5^. This bidirectional modulation of cortical activity is mediated by thalamic relay neurons whose activity is modulated by BG output nuclei and in turn project back to cortex, completing the CBGT loop^1,6^. In mice, CBGT circuits begin to form during late embryogenesis and undergo protracted maturation during postnatal stages^7–9^. While substantial research has focused on cortical activity regulation of striatal development, the influence of the BG on cortical circuit maturation remains poorly understood. However, the recurrent architecture of CBGT circuits enables tight coupling between cortical and basal ganglia activity^6,8,10,11^, suggesting a potential regulatory role of the BG in establishing early cortical connectivity.

The postnatal maturation of cortical circuits involves extensive synaptic remodeling between excitatory glutamatergic pyramidal neurons (PYRs) and local GABAergic interneurons (INs). In sensory cortices, this remodeling is strongly shaped by neural activity, with manipulations of sensory experience or thalamocortical input having pronounced effects on the maturation of cortical connectivity^12–14^. Concurrently, the maturation of GABAergic INs is critical for the emergence of structured cortical activity patterns and for regulating synaptic plasticity mechanisms essential for cortical development^15–17^. Compared to sensory areas, the activity-dependent mechanisms that guide the postnatal maturation of dmPFC circuits are less understood, partly due to the absence of direct sensory input to frontal cortical regions. Medial thalamic nuclei^18–20^ provide dense innervation to the dmPFC from the first postnatal days^9,21^, and strongly drive both PYRs and INs in adulthood^18,20^. Furthermore, these thalamic nuclei are under the modulatory control of BG output nuclei, and manipulations of their activity during adolescence alter dmPFC circuit activity and connectivity^11,22,23^. These findings suggest that BG-thalamocortical pathways may play a developmental role in the prefrontal cortex similar to the role of thalamocortical input in sensory areas. However, whether the BG modulate dmPFC activity or connectivity during early postnatal periods, and whether disruptions in BG output impair dmPFC circuit development at these stages remains unclear.

The initial wiring of CBGT circuits in mice is established perinatally, but most synaptic connections within these networks mature only after the onset of sensory experience during the second postnatal week^8,9,13,14^. At this stage, mature patterns of neural activity emerge concurrently in cortical, thalamic, and BG circuits, suggesting a coordinated developmental process^7,24,25^. These postnatal changes in network activity drive widespread remodeling of synaptic connectivity across CBGT circuits^8,26–29^. Notably, developmentally silencing either iSPNs or dSPNs bidirectionally regulates the maturation of striatal afferent connectivity ^30^, whereas chemogenetic inhibition of iSPNs during the second postnatal week leads to persistent deficits in motivated and locomotion behaviors^31^, indicating that early perturbations of striatal activity induce long-lasting structural and functional impairments. Based on this evidence, we hypothesized that dSPNs and iSPNs regulate cortical activity during early postnatal development, and that BG-mediated modulation of network dynamics contributes to cortical circuit maturation.

## Results

To investigate whether striatal circuit output modulates cortical activity during early postnatal development, we selectively ablated dSPNs or iSPNs during the first postnatal week and assessed the resulting effects on dmPFC neuronal activity using *in vivo* two-photon calcium (Ca^2+^) imaging at postnatal days (P)14–18. Leveraging the strictly ipsilateral organization of BG–thalamocortical relay pathways, we performed unilateral SPN ablations and performed within-animal comparisons between the ablated (ipsilateral) and control (contralateral) dmPFC hemispheres **(Figure 1A)**. SPN ablation was targeted to the dorsomedial striatum (DMS), which receives dense projections from the dmPFC. To ablate dSPNs or iSPNs, we injected an adeno-associated virus (AAV) expressing Cre-dependent Caspase-3 (AAV5-DIO-taCasp3-TEVp) unilaterally into the DMS of Drd1a-Cre (D1-Cre) or Adora2a-Cre (A2A-Cre) mice, respectively, at P1 (Figure 1B). To validate the efficacy and specificity of the ablation, we performed parallel injections in reporter mice carrying a floxed tdTomato allele (*Ai9*) and quantified tdTomato+ SPNs in DMS at P15. Casp3 ablation reduced dSPN density by ∼28% in D1-Cre mice and iSPN density by ∼18% in A2A-Cre mice **(Figure 1C–D)**. Because a subset of cortical neurons in D1-Cre mice express Cre, we quantified tdTomato^+^ neurons in D1-Cre x Ai9 mice within the dmPFC and sensorimotor regions along the injection track. No evidence of cortical cell loss was observed in either region, confirming that the ablation was restricted to the striatum **(Figure S1A-B).** To further assess viral targeting specificity, we injected an AAV expressing Cre-dependent GFP in A2A-Cre mice and confirmed that AAV expression was restricted to the dorsal striatum with minimal spread to ventral striatal regions **(Figure S1C)**. In addition, *in situ* hybridization in D1-Cre mice injected with AAV-DIO-Casp3 revealed a selective reduction in D1R-expressing neurons with no significant change in D2R-expressing cells, confirming that ablation was restricted to the Cre-expressing dSPN population **(Figure S1D)**. To monitor cortical activity, we bilaterally injected AAV9-hSyn-GCaMP8s in the dmPFC at P7 to express the calcium sensor GCaMP8s in neurons. At P14–P18, spontaneous Ca^2+^ transients reflecting neuronal activity were recorded simultaneously from both hemispheres in awake, head-fixed mice using two-photon microscopy **(Figure 1E** and **Supplementary Video 1)**. Traces from individual neurons were analyzed to extract Ca^2+^ event frequency, amplitude, and decay kinetics **(Figure 1F-G**; see Methods).

**Fig. 1.**
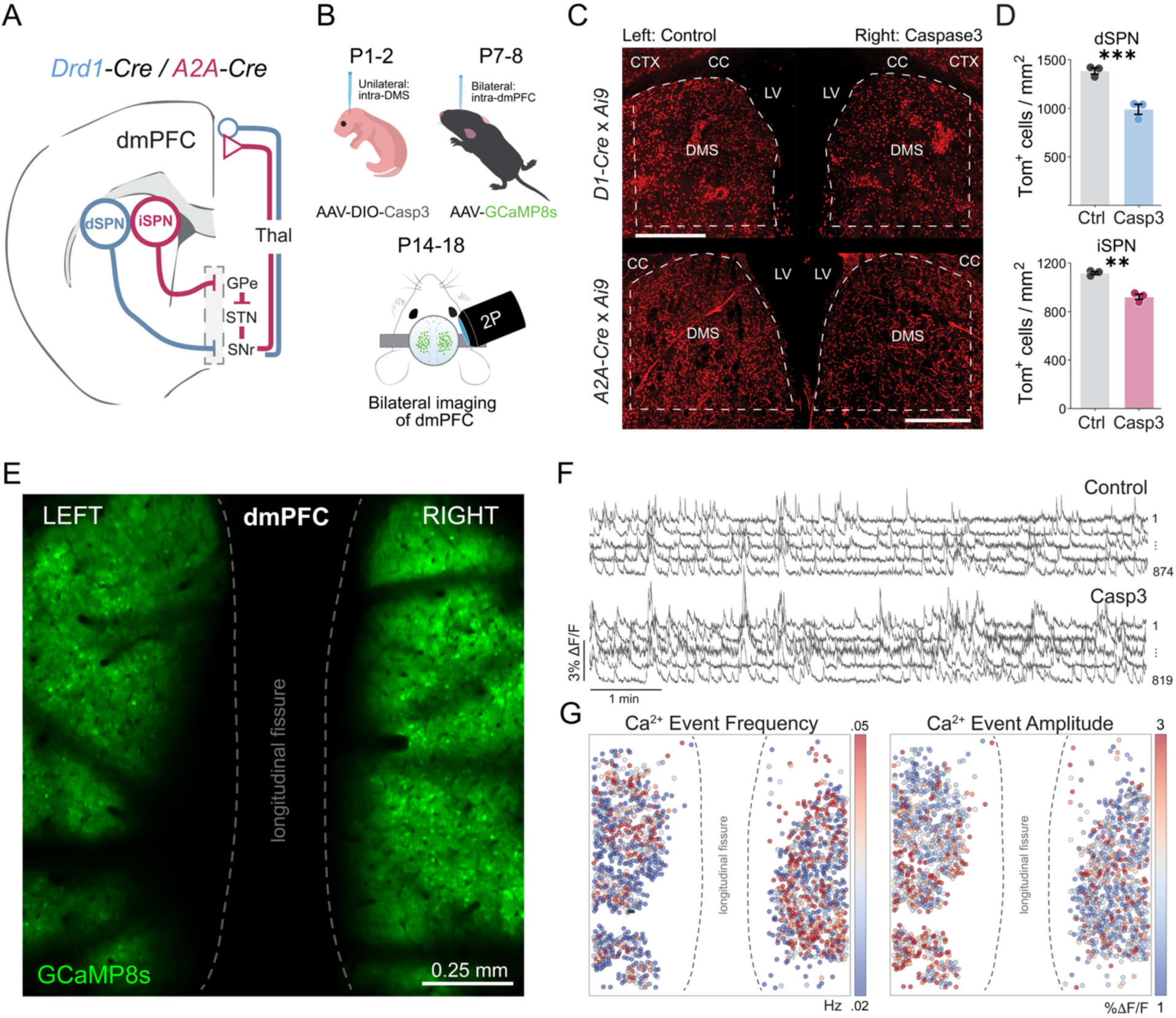
Bilateral imaging of dmPFC activity during postnatal development using two-photon microscopy. **(A)** Schematic of corticostriatal and basal ganglia loops showing direct (dSPN, blue) and indirect (iSPN, magenta) pathway neurons and their projections from the dorsomedial striatum (DMS) to downstream targets including the globus pallidus externus (GPe), subthalamic nucleus (STN), and substantia nigra pars reticulata (SNr). Cre-driver lines for dSPNs (Drd1-Cre) and iSPNs (A2A-Cre). **B)** Timeline of experimental approach. AAV-DIO-Caspase3 was injected unilaterally into the DMS of Drd1-Cre or A2A-Cre mice at P1–P2 to induce selective ablation of dSPNs or iSPNs, respectively. At P7–P8, AAV9-hSyn-GCaMP8s was bilaterally injected into the dmPFC, followed by cranial window implantation and acute two-photon imaging at P14–18 to assess cortical activity. **(C)** Representative images showing tdTomato (Tom⁺)-labeled SPNs in the DMS of control (left) and Caspase3-ablated (right) hemispheres in Drd1-Cre (top) and A2A-Cre (bottom) mice crossed with *Ai9* reporter. SPN loss is restricted to the targeted hemisphere. CTX, cortex; CC, corpus callosum; LV, lateral ventricle. Scale bars=0.5mm. **(D)** Quantification of Tom⁺ cell density in the DMS confirms that (top) ablation significantly reduced dSPNs relative to the untreated hemisphere (Control: mean±SEM = 992±52; Casp3: mean±SEM = 1386±31). Similarly, (bottom) Casp3-ablation reduced iSPN density relative to the control hemisphere (Control: mean±SEM = 1118±12; Casp3: mean±SEM = 922±22). RM-2way ANOVA detected significant main effects of Cell-Type (p=0.0168), Treatment (p<0.0001), and their interaction (p=0.0054), both dSPN (p=0.0002) and iSPN (p=0.0031) effects surviving Bonferroni corrections. Means of n=9 sections from N=3 mice/group. **(E)** Two-photon imaging of bilateral GCaMP8s expression in dmPFC of awake P15 mouse. **(F)** Representative traces of ΔF/F from GCaMP8s-expressing neurons in left and right dmPFC recorded in (E). Traces are vertically stacked and aligned by time; elevated activity is apparent in the Caspase3-ablated hemisphere. **(G)** Spatial maps of calcium event frequency (left) and amplitude (right) from dmPFC neurons in (E). Each dot represents a cell with color scaled to event frequency (Hz) or amplitude (%ΔF/F).

Postnatal ablation of iSPNs in A2A-Cre mice robustly increased the frequency of Ca^2+^ events in the Casp3-ablated hemisphere compared to the control **(Figures 2B** and **S2A)**. To assess hemispheric bias in event rates, we calculated an asymmetry index (AI) as (Casp3– Control)/(Casp3+Control) and tested its deviation from zero using a hierarchical bootstrap across recordings and mice (see Methods). The resulting 95% confidence interval excluded zero and significantly differed from shuffled controls, indicating a robust hemispheric difference.

**Fig. 2.**
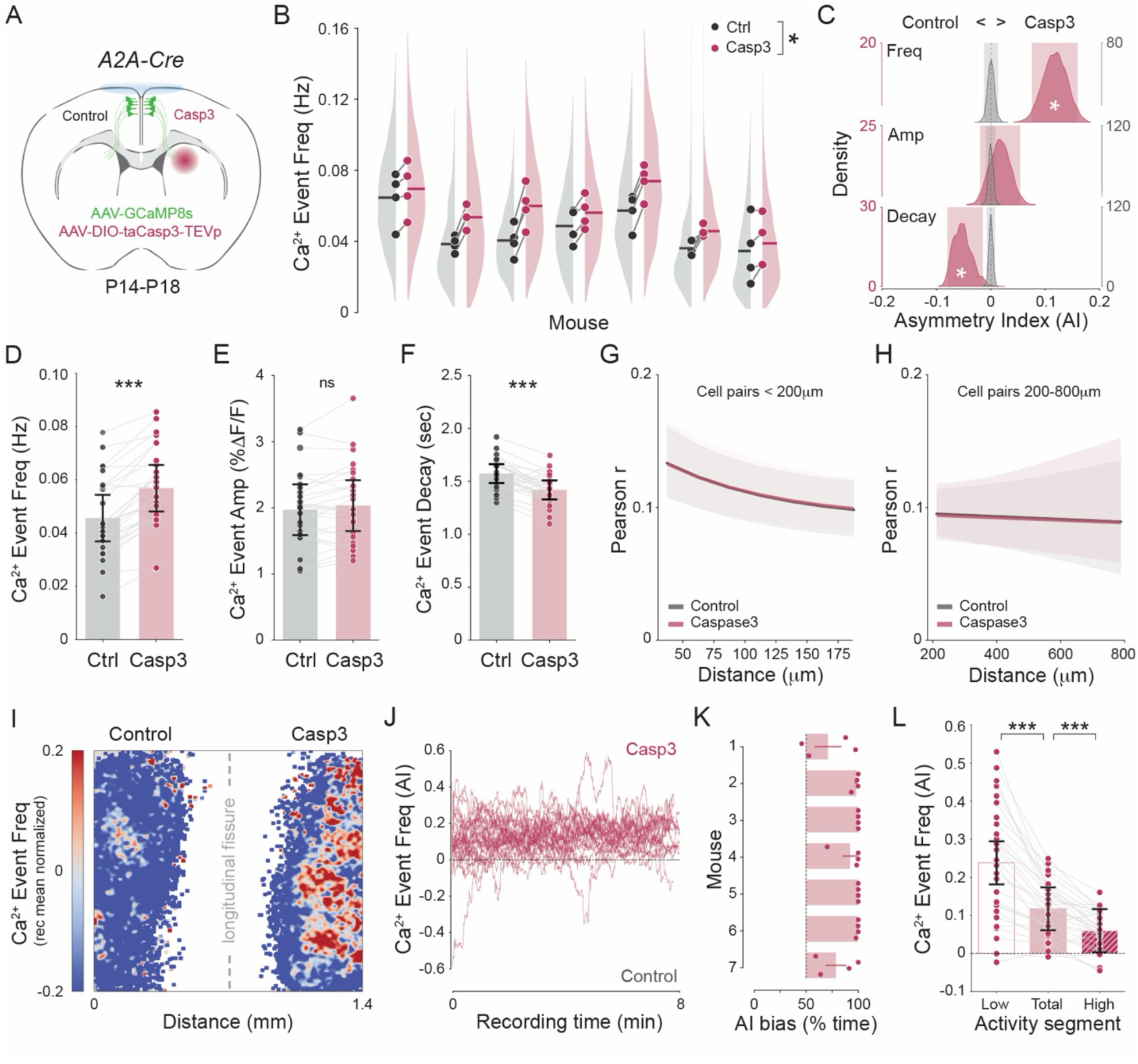
Postnatal ablation of iSPNs increases dmPFC neural activity. **(A)** Experimental schematic. Adora2a-Cre mice received unilateral AAV5-DIO-Casp3-TEVp in the DMS at P1 and bilateral AAV9-hSyn-GCaMP8s in dmPFC; two-photon imaging was performed P14–P18. **(B)** Ca^2+^ event frequency per recording in control (gray) versus Casp3-ablated (magenta) hemispheres across seven mice. Half-violin plots represent each mouse’s bootstrapped distribution of mean frequencies comprising all recordings; points represent means of individual recordings; horizontal bars indicate mouse means (Ctrl: 0.045±0.011 Hz; Casp3: 0.057±0.012 Hz; Wilcoxon signed-rank, *p*=0.016). **(C)** Hierarchical-bootstrap distributions of the asymmetry index (AI, *(Casp3–Ctrl)/(Casp3+Ctrl)*) for Ca^2+^ event frequency (top), amplitude (middle) and decay time (bottom). Magenta KDE-curves (left axis) plot true data; gray curves (right axis) plot shuffled-label control. Shaded bands denote CI_95%_. Positive values indicate larger responses in the Casp3 hemisphere (Frequency: 0.12 [0.07,0.15]; Amplitude: 0.017 [–0.019,0.054]; decay: –0.052 [–0.079,–0.015]). **(D–F)** Linear mixed-effects model estimates for frequency (D), amplitude (E) and decay time (F). Bars are fixed-effect means±CI_95%_; dots represent paired-recording means connected by light gray lines. Fixed-effect *p*-values and CI_95%_: (Frequency *β*=0.011 [0.01,0.011], *p*=1.79×10^−298^; Amplitude *β*=0.03 [–0.011,0.137], *p*=0.097; decay time *β*=–0.153 [–0.210,–0.097], *p*=8.48×10^−8^). **(G–H)** Distance-dependent Pearson *r* for proximal (<200 µm) and distal (200–800 µm) neuron pairs. Lines represent mean fits across 5,000 bootstrap iterations; shaded areas are CI_95%_. Exponential fits (<200 µm): Casp3-Ctrl decay=0.0001 [−0.004,0.005]; Δmax=-0.002 [−0.014,0.01]; Δbaseline=0.001 [−0.015,0.014]. Linear fits (200–800 µm): Casp3-Ctrl Δslope=-0.00001 [−0.00009,0.00007], Δintercept=-0.0013 [−0.026,0.032]. **(I)** Spatial heat-maps of normalized Ca^2+^ event frequency in the FOV in control (left) and Caspase3 (right) hemispheres averaged across all recordings. **(J)** Time course of the Ca^2+^ event-frequency AI of each recording. Values above zero represent increased activity in Casp3-ablated hemisphere relative to control. iSPN ablation leads to sustained positive asymmetry compared to controls. **(K)** Each animal’s percentage of recording time with asymmetry>0. Bars=mean±SEM of recordings; dots are individual recordings; dashed line at 50%. Mouse mean=98±4.8 % (N=7), Wilcoxon signed-rank, *p*=0.016. **(L)** AI calculated across network states (Low, Total, High). Bars represent LMM-estimated marginal means (Total as intercept)±CI_95%_; dots=recording means; gray lines connect recordings. Pairwise contrasts: Low vs Total and Total vs High are both p<0.001. LMM fixed-effect *p*-values [CI_95%_]: intercept=4.45×10^−5^ [0.06,0.17]; Low vs Total=4.9×10^−11^ [0.08,0.16]; High vs Total=0.00181 [–0.09,–0.02]. Random effects non-significant.

These findings were further supported by linear mixed-effects models (LMM) with hemisphere (Control vs. Casp3) as a fixed effect and mouse and recording as nested random intercepts, confirming a significant increase in event frequency in the Casp3 hemisphere. Unlike event frequency, the amplitude of Ca^2+^ events was not altered by Casp3 ablation **(Figures 2E and S2B)**. However, the event decay time was significantly reduced, suggesting potential changes in the underlying patterns of spiking activity or intracellular calcium dynamics **(Figures 2F and S2C)**. To compare the spatial correlation structure of neural activity between hemispheres, we quantified pairwise Pearson correlation coefficients (*r*) of deconvolved Ca^2+^ signal as a function of inter-neuronal distance. As expected, Pearson *r* declined substantially with increasing distance **(Figure S2D-E).** To account for this distance-dependent effect, we modeled short-range pairs (<200μm) using an exponential fit and long-range pairs (200–800μm) using a linear fit with bootstrapped estimation of confidence intervals. We observed no statistically significant differences in Pearson *r* values between hemispheres in either distance range **(Figure 2G-H)**.

To examine the spatial pattern of neural activity differences across hemispheres, we generated 2D binned maps of normalized calcium event frequency using a standardized spatial grid. On average, the effect of Casp3 ablation was broadly distributed across the anterior–posterior axis, without evidence of a spatially restricted activity changes within the recorded field of view **(Figure 2I)**. We also analyzed hemispheric differences in neural activity across time by computing the AI of event frequency throughout each recording. Notably, most recordings showed AI values consistently greater than zero, indicating that neural activity in the Casp3-ablated hemisphere was elevated relative to the control hemisphere throughout the entire recording duration **(Figure 2J)**. To quantify this effect, we calculated the proportion of time during which AI > 0 for each recording and for each mouse. This analysis revealed a robust and statistically significant hemispheric bias of increased Ca^2+^ activity in the Casp3-ablated side. To assess whether iSPN ablation-induced changes in neural activity varied across brain states, we segmented each recording into low and high activity periods and computed the AI for different Ca^2+^ event parameters within each activity segment. Across all conditions, the Casp3-ablated hemisphere consistently exhibited higher activity levels. However, there was a significant effect of activity state on event frequency, AI being most elevated during low activity periods and lower during high activity states **(Figure 2L)**. Interestingly, AI of event amplitude also increased in relation to the Control hemisphere specifically during low activity periods **(Figure S2F)**. In contrast, the asymmetry in Ca^2+^ event decay was consistent across activity periods with no differences in pairwise Pearson correlation observed in either period **(Figure S2K-P)**. These findings indicate that the increase in dmPFC activity resulting from iSPN ablation is more pronounced during quiescent network states.

Contrary to the effects observed in A2A-Cre mice, postnatal ablation of dSPNs in D1-Cre mice resulted in a significant reduction in Ca^2+^ event frequency in the Casp3-ablated hemisphere relative to the control **(Figures 3B** and **S3A)**. The bootstrapped AI for event frequency was significantly below zero and differed from shuffled distributions, indicating a robust hemispheric bias **(Figure 3C)**. LMM modeling confirmed a significant reduction in event frequency in the Casp3 hemisphere **(Figure 3D)**. In contrast to iSPN ablation, dSPN ablation also produced a significant increase in Ca^2+^ event amplitude **(Figures 3C, 3E,** and **S3B)**. Additionally, the decay time of Ca^2+^ events was modestly but significantly prolonged in the ablated hemisphere, corroborated by both LMM and mouse-level comparisons **(Figures 3F** and **S3C)**. Analysis of intra-hemispheric pairwise correlations between neurons within 200 μm revealed no significant difference between hemispheres, consistent with findings in A2A-Cre mice **(Figure 3G)**. Although we observed a trend toward reduced Pearson correlation at longer distances (200– 800μm) in the dSPN-ablated hemisphere **(Figures 3H** and **S3E)**, this effect was not consistent across mice **(Figure S3N)** and did not reach statistical significance in either hierarchical bootstrap or LMM analyses. Binned activity maps revealed a widespread reduction in activity in the dSPN-ablated hemisphere along the antero-posterior axis **(Figure 3I)**. Time-resolved AI analysis demonstrated this reduction in activity persisted throughout the recording duration in most mice, suggesting a sustained suppression of network activity **(Figure 3J)**. We further quantified the proportion of time with AI > 0, which was significantly lower than expected by chance **(Figure 3K)**. We next examined whether network state modulated hemispheric asymmetry. In contrast to iSPN ablation, which produced the greatest asymmetry during low activity periods, the reduction in event frequency following dSPN ablation was most pronounced during higher activity states **(Figure 3L)**. A small but statistically significant increase in Ca^2+^ event amplitude was also observed during low activity periods **(Figure S3F)**, whereas no state-dependent differences were found for event decay **(Figure S3G)** or for pairwise correlations **(Figure S3H–P)**. Together, these results reveal that iSPN and dSPN ablations exert opposite modulatory influences on cortical activity. While iSPN loss preferentially enhances activity during quiescent network states, dSPN loss leads to a persistent reduction in activity that is most pronounced during periods of elevated cortical activation.

**Fig. 3.**
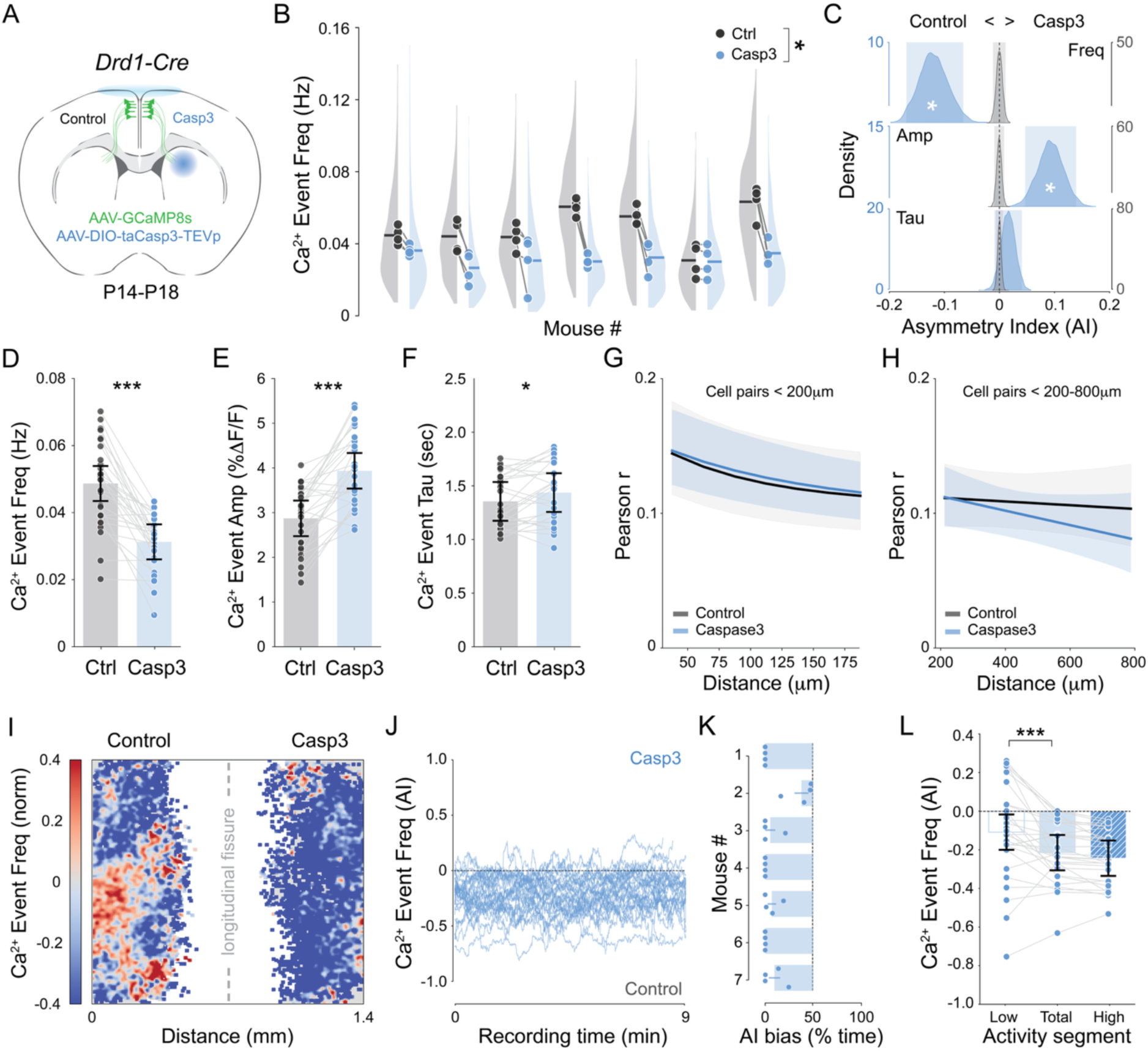
Postnatal ablation of dSPNs reduces dmPFC neural activity. **(A)** Experimental schematic. Drd1-Cre mice received unilateral AAV5-DIO-Casp3-TEVp in the DMS at P1 and bilateral AAV9-hSyn-GCaMP8s in dmPFC; two-photon imaging was performed P14–P18. **(B)** Ca^2+^ event frequency per recording in control (gray) versus Casp3-ablated (blue) hemispheres across seven mice. Half-violin plots represent each mouse’s bootstrapped distribution of mean frequencies of all recordings and points represent mean of individual recordings; horizontal bars indicate mouse averages (Ctrl: 0.047±0.011 Hz; Casp3: 0.031±0.003 Hz; Wilcoxon signed-rank, *p*=0.016). **(C)** Hierarchical-bootstrap distributions of the asymmetry index (AI, *(Casp3–Ctrl)/(Casp3+Ctrl)*) for Ca^2+^ event frequency (top), amplitude (middle) and decay time (bottom). Blue KDE-curves (left axis) plot true data; gray curves (right axis) plot shuffled-label control. Shaded bands are CI_95%_. Positive values indicate larger responses in the Casp3 hemisphere (Frequency: −0.21 [−0.30,-0.12]; Amplitude: 0.16 [0.08,0.24]; decay time: 0.027 [–0.015,0.070]). **(D–F)** LMM model estimates for frequency (D), amplitude (E) and decay time (F). Bars are fixed-effect means±CI_95%_; dots represent paired-recording means connected by light gray lines. Fixed-effect *p*-values and CI_95%_: (Frequency *β*=-0.017 [−0.022,-0.013], *p*=4.02×10^−15^; Amplitude *β*=1.065 [0.77,1.36], *p*=1.42×10^−12^; decay time *β*=0.082 [0.014,0.150], *p*=0.018). **(G–H)** Distance-dependent Pearson *r* for proximal (<200 µm) and distal (200–800 µm) neuron pairs. Lines represent mean fits across 5,000 bootstrap iterations; shaded areas are 95 % CIs. Exponential fits (<200 µm): Casp3-Ctrl Δdecay=0.0038 [−0.0041,0.012]; Δmax=0.09 [−0.015, 0.061]; Δbaseline=-0.096 [−0.074,0.016]. Linear fits (200–800 µm): Casp3-Ctrl Δslope=-0.00004 [−0.00014,0.00004], Δintercept=0.0091 [−0.026,0.048]. **(I)** Spatial heat-maps of normalized Ca^2+^ event frequency in control (left) and Caspase3 (right) hemispheres averaged across all recordings. **(J)** Time course of the Ca^2+^ event-frequency AI of each recording. Values above zero represent increased activity in Casp3-ablated hemisphere relative to control. dSPN ablation leads to sustained positive asymmetry compared to controls. **(K)** Each animal’s percentage of recording time with asymmetry>0. Bars=mean±SEM of recordings; dots are individual recordings; dashed line at 50%. Mouse mean=5.4±12.8% (n = 7), Wilcoxon signed-rank, *p*=0.016. **(L)** AI calculated across network states (Low, Total, High). Bars show LMM-estimated marginal means (Total as intercept)±CI_95%_; blue dots=recording means; gray lines connect recordings. Pairwise contrasts: Low vs Total *p*=0.00027; High vs Total *p*=0.325. LMM fixed-effect *p* values [CI_95%_]: intercept=4.66×10^−6^ [–0.31,–0.12]; Low vs Total=0.00027 [0.05,0.16]; High vs Total=0.325 [–0.09,0.03]. Random effects: Group variance, *p*=0.287 [–0.60,2.02]; Recording variance, *p*=0.0293 [0.13,2.42].

To determine whether striatal perturbations influence the maturation of synaptic connectivity in the dmPFC, we performed whole-cell voltage-clamp recordings from L2/3 pyramidal neurons in acute slices from P15 A2A-Cre and D1-Cre mice that received neonatal Caspase3 injections in the DMS **(Figure 4A)**. We measured miniature excitatory (mEPSCs) and inhibitory (mIPSCs) postsynaptic currents to assess synaptic input properties in the hemisphere ipsilateral to the manipulation. In A2A-Cre mice, ablation of iSPNs selectively reduced the frequency of GABAergic mIPSCs without altering mEPSC frequency **(Figure 4B–C)**, leading to an increase in the excitatory/inhibitory (E/I) frequency ratio **(Figure 4D–E)**. In contrast, mEPSC and mIPSC amplitudes were largely similar between groups, with only a subtle reduction in mIPSC amplitude following iSPN ablation **(Figure 4F–G)**. No changes were observed in E/I amplitude ratios **(Figure 4H)**, and there was no significant correlation between excitatory and inhibitory amplitudes within cells **(Figure 4I)**. These findings suggest that iSPN loss primarily reduces inhibitory synaptic drive onto dmPFC pyramidal neurons. In D1-Cre mice, ablation of dSPNs did not affect the frequency of inhibitory input, but mEPSC frequency was slightly increased **(Figure 4K–L)**. Although both mEPSC and mIPSC amplitudes were similar across groups **(Figure 4O–P)**, the E/I frequency ratio was modestly elevated following dSPN ablation **(Figure 4M)**. Importantly, the within-cell relationship of excitatory and inhibitory input remained intact **(Figure 4N)** and amplitude relationships were unaffected **(Figure 4Q–R)**. These results indicate that dSPN ablation does not substantially impair the maturation of cortical inhibitory synaptic input. Passive intrinsic properties and mPSC kinetics were generally preserved in both A2A-Cre and D1-Cre cohorts with the exception of a small reduction in membrane capacitance in dmPFC neurons of dSPN-ablated mice **(Figures S4A–D** and **S4M–P)**. Distributions of mPSC inter-event intervals and amplitudes supported these findings and highlighted altered inhibitory input distribution in the A2A-Cre group **(Figures S4E–H** and **S4Q–T)**. Kinetic parameters such as rise and decay times of mPSCs were comparable between groups **(Figures S4I–L** and **S4U–X)**. Collectively, these data indicate that loss of iSPNs and dSPNs during early postnatal development induces different adaptations in synaptic connectivity of dmPFC L2/3 pyramidal neurons, with iSPN loss causing a pronounced disruption of GABAergic connectivity.

**Figure 4.**
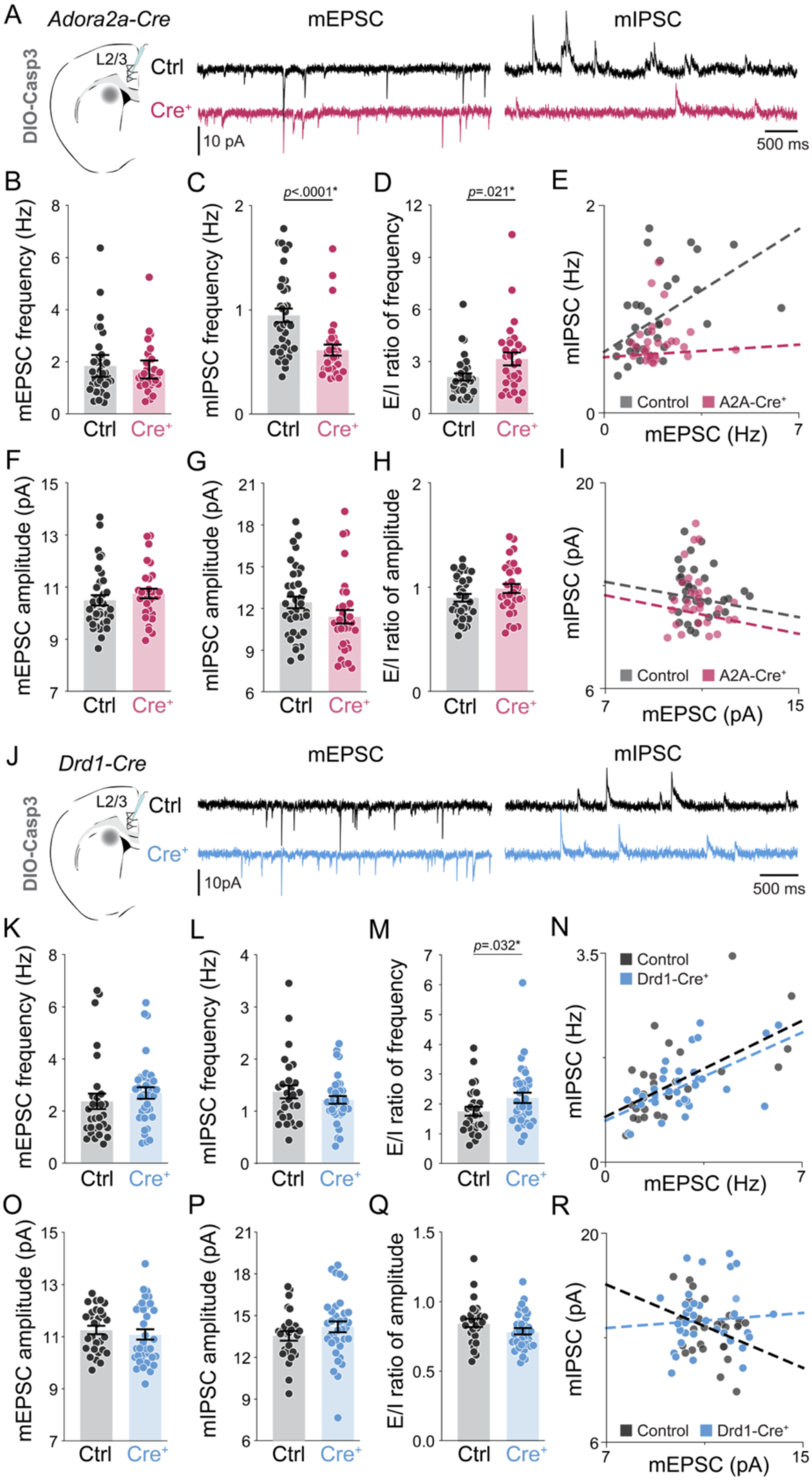
iSPN ablation reduces GABAergic input and alters E/I balance in dmPFC pyramidal neurons. **(A)** Left: Schematic of whole-cell voltage-clamp recordings from L2/3 pyramidal neurons in dmPFC of P14–P15 A2A-Cre mice injected neonatally with AAV5-DIO-Caspase3 in DMS. Right: Representative traces of AMPAR-mediated mEPSCs (V_h_=–70 mV) and GABA-mediated mIPSCs (V_h_=0 mV) from control (black) and Casp3-ablated hemispheres (magenta). Control: n=36 cells from N=4 mice. Casp3: n=30 cells from N=3 mice. **(B)** mEPSC frequency in A2A-Cre mice did not differ from controls (Control=1.84±0.206 Hz; A2A-Cre=1.70±0.175 Hz; Mann-Whitney, *p*=0.863). **(C)** mIPSC frequency was significantly reduced by iSPN ablation (Control=0.95±0.066 Hz; A2A-Cre=0.61±0.050 Hz; Mann–Whitney, *p*<0.0001). This effect was confirmed using an LMM accounting for events nested within cells and mice: model intercept inter-event interval=1240.4±91.7 ms; fixed effect of Cre^+^ was significant (*β*=583.4±135.3ms, *p*=0.000016). **(D)** E/I frequency ratio was significantly increased in A2A-Cre mice (Control=2.12±0.204; A2A-Cre=3.16±0.364; Mann-Whitney, *p*=0.021). **(E)** Robust linear regression of within-cell mEPSC vs mIPSC frequency revealed disrupted excitatory-inhibitory input relationship in A2A-Cre mice (Control slope=0.17±0.050, CI_95%_ [0.070,0.268], *p*<0.001; A2A-Cre slope=0.018±0.035, CI_95%_ [–0.051,0.086], *p*=0.615). **(F)** mEPSC amplitude was not significantly altered (Control=10.49±0.196 pA; A2A-Cre=10.75±0.187 pA; *p*=0.156). **(G)** mIPSC amplitude showed a marginal reduction in A2A-Cre mice (Control=12.43±0.426 pA; A2A-Cre=11.39±0.502 pA; *p*=0.054). **(H)** E/I amplitude ratio did not differ significantly between groups (Control=0.898±0.036; A2A-Cre=0.986±0.046; *p*=0.177). **(I)** Regression of mEPSC vs mIPSC amplitude did not yield a correlation in either group (Control slope=–0.300±0.432, CI_95%_ [–1.146,0.546], p=0.487; A2A-Cre slope =–0.328±0.454, CI_95%_ [– 1.217,0.562], *p*=0.470). **(J)** Schematic as in (A), for recordings from D1-Cre mice (Casp3-injected group depicted in blue). Control n = 29 cells from 3 mice, Casp3 n = 35 cells from 3 mice. **(K)** mEPSC frequency in D1-Cre mice was slightly elevated but did not reach significance (Control=2.37±0.308 Hz; D1-Cre=2.70±0.218 Hz, n=35 cells; *p*=0.059). **(L)** mIPSC frequency was unaffected by dSPN ablation (Control=1.37±0.125 Hz; D1-Cre=1.21±0.077 Hz; *p*=0.569). **(M)** E/I frequency ratio was significantly increased in D1-Cre mice (Control=1.75±0.148; D1-Cre=2.21±0.171; *p*=0.032). **(N)** Robust linear regression of mEPSC vs mIPSC frequency showed preserved positive E/I relationship in both groups (Control slope=0.230±0.049, CI_95%_ [0.134,0.325], *p*<0.001; D1-Cre slope=0.211±0.052, CI_95%_ [0.108,0.313], *p*<0.001). **(O)** mEPSC amplitude did not differ between groups (Control=11.26±0.157 pA; D1-Cre=11.08±0.194 pA; *p*=0.421). **(P)** mIPSC amplitude was similar across groups (Control=12.88±0.509 pA; D1-Cre=12.50±0.486 pA; *p*=0.568). **(Q)** E/I amplitude ratio was not different (Control=0.848±0.029; D1-Cre=0.788±0.022; *p*=0.102). **(R)** Regression of mEPSC vs mIPSC amplitude did not reveal a significant relationship in either group (Control slope=–0.697, CI_95%_ [–1.496,0.103], *p*=0.09; D1-Cre slope=0.126, CI_95%_= [– 0.483,0.735], *p*=0.406).

## Discussion

Our results demonstrate that imbalances in striatal output during early postnatal development alter the activity and synaptic connectivity of dmPFC circuits. Selective ablation of dSPNs and iSPNs produced opposing effects on cortical activity, consistent with prior studies in adult mice showing that direct and indirect pathway neurons exert antagonistic influence on cortical dynamics and behavior^2–5^. Our findings suggest that basal ganglia modulation of cortical activity is established during the first two postnatal weeks and align with previous work showing that postnatal silencing of dSPNs or iSPNs decreases or increases striatal glutamatergic afferent connectivity, respectively^30^. Although cortical dynamics were not directly assessed in that study, our data support a role for SPN output in modulating network activity, potentially driving widespread connectivity adaptations across CBGT circuits. The effects of striatal manipulations on dmPFC activity were observed throughout most of the recording period **(Figures 2J and 3J)**, indicating a persistent shift in cortical activity, yet the magnitude of imbalance varied with network activity state. iSPN ablation led to a marked increase in dmPFC Ca^2+^ event frequency during low-activity or quiescent periods **(Figure 2L)**, suggesting that the indirect pathway normally constrains cortical activity. In contrast, dSPN ablation caused the greatest reduction in cortical activity during high-activity periods **(Figure 3L)**, indicating a role of the direct pathway in sustaining recurrent cortical engagement when the network is active^11,22^.

Ablation of iSPNs disrupted the maturation of synaptic connectivity in L2/3 pyramidal neurons of the dmPFC **(Figure 4)**, leading to a marked reduction in mIPSC frequency and increased synaptic E/I **(Figures 4D-E)**. These unexpected findings raise the possibility that BG activity contributes to the development of cortical connectivity. BG-recipient thalamic nuclei, including the mediodorsal and ventromedial thalamus, are major drivers of dmPFC pyramidal neurons as well as diverse classes of cortical interneurons^18,19^. These thalamocortical projections reach the upper cortical plate perinatally and peak in density during the second postnatal week^9,21^. In sensory cortices, manipulations of sensory input or thalamocortical activity during early development robustly influence the maturation of cortical circuits and GABAergic interneurons^23,32–34^, suggesting that similar thalamocortical mechanisms may regulate prefrontal cortical development. Activity-dependent changes in interneuron maturation are complex and depend on cortical area, cell-type, and developmental timing^35^. In our study, iSPN ablation increased cortical activity while reducing inhibitory input, suggesting a potential non-homeostatic adaptation in which early hyperactivity—induced by abnormal BG output—disrupts or delays interneuron development or GABAergic synaptogenesis. Whether this reduction in inhibition causally contributes to cortical hyperactivity, or instead reflects a compensatory adaptation, remains unclear and will require further mechanistic investigation. In contrast, dSPN ablation did not alter mIPSC frequency, although the observed changes in membrane capacitance raise the possibility that properties not assessed in this study may have been affected^31^.

Experience-dependent plasticity is a hallmark of cortical development, enabling circuits to adapt in response to environmental inputs to promote learning and flexible behavior. While subcortical structures such as the basal ganglia are well known for their roles in reinforcement learning and action selection, whether their activity influences the organization of cortical networks remains unclear. Our findings suggest that striatal circuits, through their modulation of cortical activity, may play a broader role in shaping experience-dependent cortical plasticity. This idea is supported by recent work showing that striatal activity is necessary for the initial establishment, but not long-term maintenance, of volitional cortical control in brain-machine interface paradigms^36^. These findings are of important clinical relevance since disrupted development of striatal circuits is a common feature of several neurodevelopmental disorders such as autism spectrum disorder (ASD)^37^. Numerous ASD mouse models exhibit imbalances in dSPN and iSPN activity that are linked to behavioral deficits^37–40^. We show that early postnatal striatal dysfunction disrupts both cortical activity dynamics and the maturation of synaptic connectivity in prefrontal circuits. The elevated dmPFC activity observed during low network states following iSPN ablation is consistent with theories implicating cortical hyperexcitability and reduced signal-to-noise as central mechanisms in ASD^41,42^. Moreover, chronic chemogenetic silencing of iSPNs during the second postnatal week results in lasting impairments in motivated behavior^31^, supporting the idea that striatopallidal dysfunction and associated cortical hyperactivity during this developmental window can cause long-lasting alterations in brain connectivity and function. Disrupted modulatory control by the basal ganglia could therefore impair the development of cortical inhibitory networks, predisposing circuits to abnormal sensory integration, impaired cognitive control, and behavioral inflexibility, core features of ASD. Although our study does not establish a direct causal relationship between these phenotypes, our findings suggest they may emerge from convergent disruptions in early circuit development. Future studies will be needed to identify the specific changes in cortical connectivity driven by early abnormal striatal activity and to determine how these alterations contribute to disease-relevant behavioral phenotypes.

## Limitations of the study

This study provides new evidence that early striatal output modulates cortical development, but several limitations should be acknowledged. First, while our manipulations were restricted to the DMS and recordings to the dmPFC, we cannot exclude the possibility that unilateral perturbations may influence contralateral cortical activity via callosal projections or induce plasticity in other brain regions. Although we observed bidirectional effects on dmPFC activity consistent with current models of basal ganglia function, we did not directly assess activity in basal ganglia output structures or thalamic relay nuclei and thus cannot definitively map the full circuit responsible for the observed changes. Second, our use of Caspase3–mediated cell ablation provides a focal and effective strategy to manipulate SPNs during early development but has limited physiological relevance. Nonetheless, it serves as a valuable first step for establishing causality and probing basal ganglia contributions to cortical circuit maturation. Third, while we observed a consistent reduction in inhibitory input following iSPN ablation, the underlying mechanism remains unknown. Future studies will be needed to map the projection- and cell-type–specific changes that contribute to altered maturation of cortical inhibition. Last, our experiments were confined to early postnatal time points, and we did not assess chronic behavioral or circuit-level consequences. Future work using temporally controlled or reversible manipulations will be essential to distinguish transient developmental effects from persistent circuit dysfunction or compensatory adaptations.

## Acknowledgements

We thank Sidney Dawkins, Lan Chen and Tasha Merchant for technical assistance and management of the mouse colony. We thank Gary Thomas for access to his Nikon ECLIPSE Ti2 confocal microscope. We are thankful to Joe Stujenske, Isabel H Bleimeister, and Michael J Leone for feedback on the manuscript. We thank Susanne Ahmari for providing the Adora2A-Cre and Drd1-Cre transgenic mice, and we thank Muhammad Nazir for his assistance with 2-photon microscope maintenance and troubleshooting. This project was supported by R.P. grant R01MH124695 and a Simons Foundation Bridge to Independence award. K.D.K. was supported by K01MH128763 and the Brain and Behavior Research Foundation (30823; P&S Fund), S.D.S. was supported by R01EY033385.

## Author Contributions

R.T.P. conceived the study and supervised the experiments. M.J. collected 2-photon imaging data; R.T.P wrote the 2P imaging analysis scripts and analyzed the data with M.J.; T.D. developed surgical protocols and performed pilot imaging experiments; M.J. and Y.S. collected whole-cell electrophysiology data; M.J. analyzed the electrophysiology data; M.M. and A.D.A. performed mouse genotyping and intracranial viral injections; M.S.P. and K.D.K. performed *in situ* hybridizations and V.V. and S.S. analyzed the in situ hybridization data; R.T.P, M.J. and T.D. wrote the manuscript with input from all the other authors.

## Declaration of generative AI and AI-assisted technologies in the writing process

During the preparation of this work the authors used ChatGPT to assist with grammar correction and grammar refinement. After using this tool, the authors reviewed and edited the content as needed and take full responsibility for the content of the publication.

## Methods

### Animals

All procedures were conducted in accordance with protocols approved by the University of Pittsburgh Institutional Animal Care and Use Committee (IACUC) and adhered to the Guide for the Care and Use of Laboratory Animals published by the National Institutes of Health. Naive C57BL/6J (JAX 000664) or Ai9 reporter (JAX 007909) mice were mated with transgenic mice expressing Cre Recombinase under the control of either Dopamine receptor type-1 promoter (D1-Cre; *Tg(Drd1a-cre)EY262Gsat/Mmcd*) or the Adenosine-2a receptor promoter (A2A-cre; *Tg(Adora2acre)KG139Gsat*)^43^. Same-age littermate comparisons were performed when possible and litter size was capped at eight pups to minimize variability in developmental rates. Pregnant females were closely monitored and neonates were considered P0 on the day of their birth. Mice were housed in a facility maintained at constant temperature and 12/12h light/dark cycle and had *ad libitum* access to water and food. All the experiments were conducted during the light phase of the cycle.

### AAV viruses

Adeno-associated viruses used in this study are commercially available (Addgene, MA, USA) AAV9-hSyn-jGCaMP8s-WPRE (Addgene #162379), and pAAV5-DIO-taCasp3-TEVp (#45580, lot #143244). For estimating the spread of virus as P1, pAAV-synP.DIO.EGFP.WPRE.hGH was used (Addgene #100043-AAV9). Viruses were aliquoted and stored at −80°C. GCaMP8s and Casp3 construct titers were ≥ 1×10¹³ and ≥ 7×10¹² GC/mL, respectively.

### Stereotaxic virus injection

D1-Cre and A2A-Cre pups were genotyped, toe-clipped, and at P1-2 placed in a heated bucket. Anesthetic hypothermia was induced by placing a pup onto ice-cold aluminum for 8-10 minutes, then transferred onto a chilled stereotaxic neonatal adaptor platform (Stoelting, #51625). The platform was kept cold by sublimating dry ice pellets in 100% ethanol. Pups were carefully inserted into the snout holder and splayed, such that their limbs were not interfering with their head positioning. Soft rubber head bars secured the head in place while the snout was leveled front-to-back and mediolaterally, such that the top of the head was ∼0.5 mm superior to Lambda, which served as a reference point. Glass pipettes (Drummond) were hand-pulled on a pipette puller (Sutter Instruments, P-1000) and broken open to achieve tip diameter of 15-25 µm, confirmed by visual inspection in a microforge (MF2, Narishige). Tubing back-filled with light mineral oil was coupled to a Hamilton syringe actuated by an injection pump (Harvard Apparatus, PHD Ultra). Injection pipettes were backfilled with light mineral oil using microfil (WPI, MF28G67-5) and through negative pressure filled with virus under dissection microscope control. To target dorsomedial striatum, the tip of the pipette was zeroed at Lambda, relative to which it was placed at AP: +2.2 and ML: ±1.0 mm. A bent G26 needle tip gently punctured the skin/skull at the pipette location, making way for the pipette tip to be lowered. The DV was offset at brain level, once the pipette tip penetrated the bone, or ∼-0.65 mm below the skin surface when the pipette tip passed through the skull without resistance or change in angle. DIO-Casp3 was delivered unilaterally at two depths, −1.9 and −1.7 mm, injecting 200 nL per depth at a rate of 50-75 nL/min. The oil-virus meniscus was marked before each injection and served to visually confirm successful injection. The pipette was left in place for ∼2 minutes after each injection to allow for diffusion. The entire duration of the neonatal injection was kept under 20 minutes to facilitate recovery. Pups were warmed up on a heating pad and then kept in a bucket until it was possible for the entire litter to return to the home cage. To aid their wellbeing, pups were mixed together with the soiled bedding and carefully placed outside of the nest, facilitating retrieval by the dam. To minimize cannibalization, breeders were fed reproductive diet (Love Mash, Bio-Serv) and were left undisturbed until pups reached P7.

At P7-8, pups used for 2P imaging were bilaterally injected with AAV9-hSyn-GCaMP8s in the dmPFC. First, pups were weighed and only underwent surgery if at least 3 grams. Anesthesia was induced in a small chamber using 4-5% isoflurane until observing 1 breath/sec and for no more than 2 minutes. Pups were placed into a neonatal nosecone (Kopf) and gently secured with Zygoma ear cups, enabling mediolateral leveling. Isoflurane concentration was gradually reduced to 1.5-2% to maintain deep anesthesia. Meanwhile, head fur was removed with Nair and disinfected using 70% ethanol. Carprofen diluted in warm sterile saline was administered subcutaneously to reduce inflammation. Scalpel blade size #11 was used to make an incision in the skin above the Bregma, which was extended rostro-caudally using surgical scissors. Connective tissue was removed using cotton Q-tips and a blunt probe (FST). Once the bone was dry, Bregma was revealed by gently pressing down on the bone plates, approximately 2.8-3.2 mm anterior to Lambda. Pipette tip was zeroed just anterior to Bregma and moved to the desired injection coordinates at AP: +0.7 to +0.8 mm and ML: ±0.3 mm, avoiding any blood vessels. At this location, a bent G26 needle tip was used to gently puncture the skull. Cold sterile saline was applied to clear any bleeding and debris, and only then was pipette tip lowered, confirming unobstructed entry into the brain tissue. At DV −0.3 mm relative to bone, pipette tip was not visible and the DV measurement was offset relative to brain surface. Within each hemisphere, AAV-GCaMP8s was delivered at two depths, starting at DV −1.0 mm and followed by a second injection at −0.75 mm. Each injection delivered 200 nL at a rate of 100-75 nL/min. Injection delivery was confirmed visually by monitoring the meniscus and any potential bleeding, which could easily obstruct the pipette. Failure to visually confirm ejection of virus from pipette was followed up by inspecting the pipette or replacing it and repeating the injection(s). Virus was allowed to diffuse for 2-4 minutes after each injection. Pipette was then withdrawn in small increments until reaching DV ∼-0.5 mm, at which point additional 50 nL were injected at a slow rate of 25 nL/min to reduce backflow. Bone surface was cleaned and dried following injections to reduce inflammation that could disrupt later procedures. A small 6-0 suture (Sharpoint) was used to close the skin incision. The entire procedure took approximately 60 minutes. Pups were allowed to recover on a heating pad until mobile, at which point the entire litter was returned outside of the nest in the home cage. Post-op carprofen or rimadyl were administered for the next three days during which pups were closely monitored. In rare cases, re-suturing or applying skin tissue adhesive (Histoacryl Blue) was required.

### Stereotaxic window surgery

At P14-18, pups expressing GCaMP8s underwent cranial window surgery to acutely measure prefrontal activity. Pups were weighed, and any pups below 6 grams or exhibiting open wounds were excluded from the procedure. Anesthesia was induced using 4% isoflurane as before and maintained at 1-1.5% once in the stereotaxic apparatus. At this age, firm non-rupture ear bars (Kopf) were used to secure the head in place just in front of the ear canal, aiming at the squamosal indent. Eyes were lubricated using an ointment (Systane) to prevent eye damage and carprofen in warm sterile saline was administered subcutaneously at the start of the procedure. Next, regrown fur was removed using Nair, and scalp was carefully cleaned with 70% ethanol. An incision was made across the skull, and skin was removed to reveal the cranial surface. The bone was scraped clean using the blunt edge of the scalpel blade and cleaned further with ice-cold sterile saline. Once the bone was dried with Q-tips, the placement and evenness of GCaMP expression were visually assessed using a blue LED and filter glasses (NightSea). Animals with unilateral or highly uneven expression of GCaMP were euthanized. While GCaMP expression was considered, craniotomy was performed starting at ∼0.5 to 1 mm anterior of Bregma and extending for ∼2.5 mm or until reaching the heavily vascularized area of the anterior sinus. The rectangular craniotomy laterally extended ±1.5-2 mm and was gently drilled with a 0.025 mm carbide drill bit powered by a foot-operated drill (Foredom). Care was taken not to penetrate the dura mater, and to stem any bleeding by alternating ice-cold sterile saline application and ice-cold 0.01% epinephrine. In case of bleeding, gelatin hemosponge (GDT) was soaked in saline and applied to the source of bleeding. With the rectangular craniotomy drilled, two Dumont number 5 forceps (FST) were inserted diagonally underneath the bone to simultaneously lift the left and the right bone plates away from the brain without compromising the midsagittal sinus. If the subsequent bleeding could not be stemmed, then the surgery was aborted. Frequent application of ice-cold sterile saline and hemosponge typically revealed access to the prefrontal cortex, which required further cleaning of blood and debris that might obscure optical access. Whenever possible, dura was not removed, but in some cases, dura had to be removed with fine forceps over both hemispheres. The brain surface was kept hydrated with sterile saline, and a glass coverslip (⌀5 mm, Electron Microscopy Sciences) was gently laid over the entire craniotomy. The edges of the coverslip overhanging the craniotomy were glued to the bone underneath (Gorilla glue). Once the saline under the coverslip was entirely confined by the glue and the coverslip, the rest of the cranial surface was scraped clean and dried so that a custom-made titanium headpost could be fitted around the cranial window. The headpost was secured using acrylic glue and leveled mediolaterally using a stereotaxic post to keep it parallel to the coverslip. The space between the headpost and the skull was filled first with acrylic glue and subsequently with 1-2 layers of black dental cement (Orthojet) to form a water-proof “well” around the cranial window. Animals received a s.c. injection of warm sterile saline (0.5 mL) and were transferred onto a heat pad, where they recovered for at least 2 hours.

### Two-photon imaging

Recovered pups aged P14-18 were head-fixed atop a heating pad in the dark and angle-corrected using a goniometer to ensure evenness of bilateral imaging. The surface of the coverslip was gently cleaned using 70% ethanol-dabbed Q-tip. Images were acquired using a resonant scanning two-photon microscope (Ultima 2P plus, Bruker, WI) at 8 Hz frame rate and 1024×1024 pixel resolution through a 16x water immersion lens (Nikon, 16x 0.8NA). dmPFC was imaged at a depth between 180μm and 320μm (zeroed at the pial surface), at the level of cortical layer 2/3. Excitation light was provided by tunable femtosecond infrared source (920 nm) wavelength laser (Insight X3, Spectra-Physics, CA). Tunable wavelength beams were combined with a dichroic mirror (ZT1040dcrb-UF3, Chroma, VT) before being routed to the microscope’s galvanometers. PrairieView software (vX5.5 Bruker, WI) was used to control the microscope. Field of view (0.8x) comprising ∼1.4 mm^2^ was centered around AP: +0.75 to +2.75 mm, relative to Bregma. Four imaging sessions were acquired from each cranial window at varying depths spaced at least 30 μm apart to capture non-overlapping populations. Images were collected while mice were alert and head restrained on top of a heat pad set to 37°C. Individual recording sessions lasted for 10 minutes. Images of GCaMP8s fluorescence collected with 920 nm excitation were processed offline in Suite2p (v 0.10.3, Cellpose 0.7.3). Two-photon calcium imaging data were preprocessed using a custom Python pipeline. Fluorescence traces (F, Fneu) and associated metadata were loaded from Suite2p-processed npy files for each recording. Neuropil-corrected fluorescence signals were computed as F_corrected_=F−0.7*F_neu_. Movement-related artifacts were identified using the mean xy displacement for every frame. Frames with xy displacement > 2 pixels as well as the surrounding 45 frames (approximately ±2.8 seconds), were excluded from further analysis; traces were concatenated and filtered with a Savitzky-Golay filter around cropping points. Fluorescence traces were converted to relative fluorescence ΔF/F signals with baseline fluorescence (F₀) estimated as the 8th percentile of values within a symmetric rolling window of 30 seconds. To remove high-frequency noise, ΔF/F traces were filtered with a wavelet denoising procedure with a Daubechies-4 (db4) wavelet at level 2. The denoised trace was reconstructed and returned alongside the noise residual. To extract calcium transients from denoised ΔF/F traces, we used the *scipy.find_peaks* function. The prominence threshold for each ROI was set to twice its baseline noise size, calculated from the interquantile range (90th–10th percentile) of the residual noise trace. Peaks were detected with the following parameters: minimum peak distance = 1 second, prominence ≥ 0.5, and width ≥ 3 frames. Onset points were defined by searching backward from each peak for a local minimum followed by a steep slope leading to the peak. Decay times were determined by identifying the time point where the smoothed trace first dropped to 37% of the peak amplitude above the onset level. Decay search windows were bounded by the onset of the next peak to avoid overlapping events. Quality control was applied to exclude non-physiological traces. Cells were retained in the dataset only if they met a multi-parameter set of criteria: minimum ΔF/F peak amplitude > 0.85, positive values for event rise time, duration, AUC, and amplitude; decay time < 3.1 s; noise < 2.5; average and minimum ΔF/F above 0; and Fneu/F ratio < 2. Traces exceeding a maximum ΔF/F of 30 were discarded. In total, 15,032 ROIs recorded in D1-Cre mice and 27,124 ROIs in A2A-Cre mice passed quality check criteria. Each D1-Cre recording comprised on average (mean ± SD) 536.8 ± 163.5 and each A2A-Cre recording 968.7 ± 433.6 cells/recording. To examine how local network dynamics influenced single-cell activity, we segmented each recording into high- and low-activity network states based on population-level event frequency. This analysis was restricted to quality-controlled ROIs located on the left hemisphere, serving as an internal control. We first computed the instantaneous event frequency by summing the number of detected ΔF/F peaks across all left-hemisphere ROIs for each frame. This time series was smoothed using a centered rolling average (2-second window) to produce a continuous measure of population activity. High-activity segments were defined as contiguous periods in which the smoothed event frequency exceeded the 75th percentile of the distribution, while low-activity segments fell below the 25th percentile. Only segments lasting at least 2 seconds were retained. To quantify how single-cell activity varied across network states, we computed event-specific properties for all detected peaks. For each ROI, peaks were assigned to high- or low-activity segments based on their temporal location. Peak frequencies were also computed separately for each state by normalizing the number of events by the cumulative duration of each activity period. For calculations of correlation coefficients, we processed ΔF/F traces using fast non-negative deconvolution using the FOOPSI algorithm implemented in the OASIS package. For each ROI, the ΔF/F trace was deconvolved using an autoregressive (AR1) model with a sparsity penalty. Event amplitude threshold of 0.05 units was used to exclude low-amplitude fluctuations and event frequencies.

### In situ hybridization

RNAscope *in situ* hybridization was performed on fresh frozen striatal tissue sections collected from D1-Cre or A2A-Cre mice injected into one hemisphere of the striatum with AAV5-DIO-taCasp3-TEVp. Brains were sectioned in a cryostat at 14um depth to isolate DMS containing sections. The left hemisphere was fiducially labeled to identify the injected hemisphere during image processing. Tissue sections were mounted onto slides and processed according to the RNAscope Multiplex Fluorescent Reagent Kit v2 protocol (UM 323100) using the following probes: D1 (RNAscope™ Probe-Mm-Drd1, cat# 461901) and D2 (RNAscope™ Probe-Mm-Drd2-C2, cat# 406501-C2). Sections were imaged at 20x magnification using an Olympus SLIDEVIEW VS200 research slide scanner. TIFF files were analyzed in the open-source software QuPath (version 0.4.4), where two GeoJSON objects encompassing the left and right striatum were traced and used on all of the sections; DAPI, FITC and CY5 channels were then isolated from these specific regions. The total number of nuclei per striatum was obtained by manually counting the DAPI channel using the count tool in Adobe Photoshop, while the FITC and Cy5 signals were segmented on QuPath by using the “dsb2018_heavy_augment.pb” pretrained model of the StarDist extension. The resulting masks were then saved as rendered SVG files, converted into JPG files, and overlapped with the DAPI channel. In order to avoid potential background segmentation, only nuclei containing masks for each respective channel were manually counted and considered positive for each one of them. Ratios of Dr1d (dSPN) to Dr2d (iSPN) positive cells were calculated and compared between injected and non-injected hemispheres to quantify SPN loss in the striatum.

### Whole-cell electrophysiology

Mice were anesthetized with isoflurane and transcardially perfused with ∼5 mL of ice-cold artificial cerebrospinal fluid (ACSF, in mM): 25 NaHCO_3_, 125 NaCl, 1.25 NaH_2_PO_4_ monohydrate, 2.5 KCl, 12.5 D-glucose, 1 MgCl_2_ hexahydrate, and 2 CaCl_2_ (on average 307.3 mOsm/kg). Upon decapitation, brains were extracted and placed on an ACSF-soaked filter paper, cerebellum was severed, and hemispheres were carefully bisected with a razor blade. The exposed posterior surface was glued to a magnetic chuck and secured in the cutting chamber containing ice-cold ACSF continuously equilibrated with carbogen (95% O_2_/5% CO_2_) and surrounded by ice. Sections containing dmPFC were cut at 275 µm (0.12 mm/s) using a vibratome (VT 2000, Leica) and incubated for 10-15 min at 32°C in a choline chloride solution containing (in mM): 25 NaHCO_3_, 1.25 NaH_2_PO_4_ monohydrate, 12.5 D-glucose, 2.5 KCl, 7 MgCl_2_ hexahydrate, 0.5 CaCl_2,_ 110 choline chloride, 11.6 ascorbic acid, and 3.1 pyruvic acid. Sections recovered for 45 min in ACSF preheated to 32°C and gradually returned to 20-22°C. Individual sections were transferred to a recording chamber attached to an upright microscope (Scientifica) and continuously perfused at 2-2.5 mL/min with equilibrated ACSF at 20-22°C. Pump-recirculated ACSF bath contained 10mM RS-CPP to abolish NMDAR-mediated currents and 1mM TTX to prevent voltage dependent sodium channel currents. To patch cells, we pulled (P-1000, 2.5×2.5 mm box filament, Sutter Instruments) pipettes from thick-walled, filamented borosilicate glass (OD 1.5 mm, ID 0.86 mm, Sutter Instruments) with resistance 2.5-3.5 MΩ when filled with the internal solution and positive pressure. To facilitate space-clamp, we filled pipettes with a cesium methanesulfonate-based solution containing (in mM): 130 CsMeSO_3_, 10 HEPES, 1.8 MgCl_2_ hydroxide, 8 sodium phosphocreatine, 10 CsCl, 3.3 QX 314 (Cl^−^ salt), 4 disodium ATP, and 0.3 sodium GTP (pH adjusted to 7.3 with CsOH, 299 mOsm/kg). Basic morphology was visualized by adding 0.7 µL of Alexa Fluor 488 hydrazide (Thermo Fisher, A10436) dissolved in water (1:100) to the internal solution that was kept on ice and dispensed through a microfil tip coupled to a 0.22 µm filter (Millipore). We targeted L2/3 cells located 150-250 µm from the medial surface, and upon forming a 2-3 GΩ seal, broke into whole-cell configuration, immediately recording passive properties. Cells were voltage clamped at V_h_ = −70mV and perfused with internal solution for 4 minutes after which mEPSCs were acquired for 5 minutes. At the end of each 10 second sweep, a test pulse injection continuously monitored input (R_in_) and series (R_s_) resistance. In order to assay inhibitory currents mediated by GABA_A_/glycine receptors, the same cell was clamped at V_h_ = 0mV, allowed to stabilize for 1 minute, and mIPSCs were acquired for 5 minutes. Currents were amplified by MultiClamp 700B (Molecular Devices), low-pass Bessel filtered at 3 kHz, digitized at 10 kHz, and acquired in ScanImage program running in MATLAB 2019a (Mathworks) and relayed by TeleConverter. Each cell recording was concluded by noting the final Rs and the presence of putative spines and an apical dendrite were confirmed by imaging (470nm, pE-300, CoolLED) the intracellular Alexa Fluor 488 signal. A 4x DIC image of the cell location in the section was obtained to confirm location in ipsilateral L2/3 dmPFC. Cells were included in analysis if they exhibited 1) initial R_s_ < 20 MΩ, 2) final R_s_ < 25 MΩ, 3) R_in_ > 100 MΩ, 4) V_h_ = −70 or 0 mV requiring < 100 pA, 5) fluorescently labeled putative spines, 6) a distinct apical dendrite, and 7) excitatory inputs decaying to tau in > 4 ms.

### Two-photon imaging data analysis

To quantify hemispheric differences in neural activity, we applied a hierarchical bootstrap procedure that estimates the asymmetry index (AI) as the ratio of (R−L)/(R+L) across multiple levels of our dataset (mouse, recording, cell). This non-parametric approach accounts for the nested structure of the data and allows direct inference on the distribution of the group-level effect without assuming normality. For each of 5000 bootstrap iterations, we resampled mice, recordings, and cells with replacement, computing a single asymmetry index per iteration from the averaged values of matched left and right cells per recording. This approach yielded a bootstrap distribution of the population-level asymmetry effect from which we derived a 95% confidence interval (CI_95%_). Significance was established if CI excluded zero. To complement this analysis, we used a linear mixed-effects model (LMM) with fixed effects for hemisphere and random intercepts for mouse and recording. To compare how correlation strength varies with distance across hemispheres, we used a hierarchical bootstrap approach to fit linear models to the distance-correlation relationship separately for left and right hemispheres. This analysis was restricted to ROI pairs within a specified distance range (<200µm and 200–800µm). For each bootstrap iteration, mice were sampled with replacement, followed by resampling of recordings and then individual ROI pairs within each recording. Binned average correlation values were computed for each side, and a linear regression was fit to these bin means. The slope and intercept of the linear fit were extracted for each hemisphere, and the fitted values across bins were stored. The process was repeated across 5000 iterations to obtain distributions of slope and intercept values for both hemispheres. From these, we calculated the average slope and intercept, confidence intervals, and the mean difference between sides. This method allowed us to quantify whether the relationship between anatomical distance and functional correlation differed systematically between hemispheres while accounting for the hierarchical structure of the data and variability across animals and recordings. To quantify the spatial patterns of neural activity differences across hemispheres, we computed two-dimensional binned activity maps from calcium imaging data. For each recording, event frequency values of both hemispheres were transformed as log2 (Hz value/ recording mean) to normalize hemispheric differences across animals and recordings. Centroid xy coordinates were extracted for each ROI within the imaging field and expanded into a 3×3 grid in both x and y dimensions. These expanded coordinates were then used to compute a binned 2D histogram of event frequency with bin counts fixed across all recordings. Gaussian smoothing (σ = 1) was applied to the group-averaged map to reduce high-frequency noise and enhance spatial structure. To assess hemispheric differences in neural activity over time, we first computed smoothed calcium event frequency traces for each recording and hemisphere using a centered moving average with a 30-second window (240 frames at 8 Hz). For each neuron, event indices were used to generate time-resolved frequency signals, normalized by cell count. These per-cell signals were then averaged and smoothed to generate per-recording traces of population activity. For each recording, we computed AI of population activity at each time point and quantified the proportion of time during which AI > 0, representing elevated activity in the Casp3-ablated hemisphere. To assess significance across animals, we tested whether the percentage of time for which AI > 0 differed from a null hypothesis of 50% (i.e., even split between hemispheres) using a Wilcoxon signed-rank test.

### Quantification of Casp3-mediated ablation

To quantify the extent of DIO-Caspase3-mediated neonatal ablation, we crossed D1-Cre and A2A-Cre mice with Ai9-tdTomato^fl/fl^ reporter mice. Offspring who inherited both Cre and a floxed Ai9 allele would therefore express tdTomato signal localized in cells containing Cre. Such pups emit a red fluorescent signal in response to green LED excitation when viewed with filter glasses (NightSea). Cre x Ai9^fl/wt^ pups were injected unilaterally at P1 with AAV-DIO-Caspase3 targeted to the DMS exactly as before (AP: +2.2; ML: ±1.0; DV: −1.9 and −1.7 mm) to mimic ablations in imaging and electrophysiology experiments. Two weeks later, brains were collected at P15-16 following a transcardial perfusion with ice-cold PBS and ice-cold 4% PFA. Following overnight post-fixation in 4% PFA at 4°C, brains were transferred to PBS and sectioned (50 µm) on a vibratome (Leica VS1200). Free-floating sections were incubated at RT in DAPI (1:5000) and mounted onto charged Superfrost (Fisher) slides, then rehydrated and coverslipped using ProLong Diamond Antifade mountant and left to cure at RT in the dark. Slide scans of samples were taken using a slide scan microscope (Olympus VS200) to select samples for further imaging. Confocal images were acquired on an inverted microscope (Nikon ECLIPSE Ti2) in the resonant galvo scan mode using a 40x water-immersion objective (Nikon). Z-stacks were acquired at 2-µm intervals and tiled (10% optimal overlap) over a large area containing the DMS and consisting of 6×8 FOVs or more. Scans were binned and averaged 4x and denoised in real time to enhance SNR. Digital zoom was set to 1x, pinhole size to 1.2 AU, and pixel size was 0.31 µm. Excitation was delivered using 405 nm and 561 nm lasers set to ∼2.7% and ∼1.75% power, to excite DAPI and tdTomato, respectively. HV/GaAsP were set to ∼85 and ∼25, respectively, with D1-Cre x Ai9 cortical cells generally being dimmer and thus requiring increased gain ∼35. Each hemisphere was imaged separately, but imaging conditions were maintained L-R, and the presence of a razor mark in the ventral surface indicated the control hemisphere. Raw z-stacks were loaded into FIJI/ImageJ to construct a maximum intensity projection using the middle ∼4 frames (8 µm), thereby ensuring even distribution of signal across the entire image. Z-projections contained the same tissue thickness in L and R hemispheres and were saved separately as TIFs. Once loaded into QuPath, a closed polygon annotation was drawn along the corpus callosum and the edge of the lateral ventricle, connecting ventrolaterally at a right angle and comprising on average 1.1-1.2 mm^2^ of dorsomedial striatum. The same polygon was transferred to the other hemisphere and mirror-reflected using a custom script. Automated cell detection was performed within each polygon annotation using the mean cell intensity of the tdTomato channel [Pixel size= 0, Background radius=3; using opening by reconstruction; Median filter radius=3; Sigma=3; Min. area=50; Max. area=200; Intensity threshold=50; Cell expansion=1 µm]. Tom^+^ cells were thresholded at 750. While no ROIs were manually added or removed at this step, we observed endothelial Tom^+^ expression in A2A-Cre tissue. To prevent contaminating neuronal cell counts with vasculature, we excluded any ROI with circularity < 0.8, thus filtering out elongated cells (18.43 ± 0.24 µm) and only keeping spherical cells (14.24 ± 0.23 µm; mean caliper length ± SD). This step, which was skipped in the D1-Cre analysis, potentially reduced measured cell density in the A2A-Cre dataset relative to the D1-Cre dataset. Cell density is expressed as the number of Tom^+^ cells divided by the total area in which they were detected. For cell quantification in dmPFC L2/3, we used only D1-Cre sections exhibiting a robust ablation in the DMS. Annotations were manually drawn in QuPath similar as above but containing 250-300 µm of tissue extending mediolaterally over the full extent of dorsal and ventral anterior cingulate cortex, corresponding to the area sampled in slice electrophysiology recordings. L1 was excluded from these annotations as it contained few to no Tom^+^ cells and because it was not targeted in whole-cell recordings. Maximum intensity projections across ∼4 optical slices were collapsed to ensure uniform sampling. Tom^+^ cells were thresholded at 150. The mean area sampled in L2/3 dmPFC consisted of ∼0.45 mm^2^. To further test the possibility that Casp3 injection leaked dorsally along the injection track or was misplaced and induced extra-striatal ablation, we quantified Tom^+^ cell density in deep L6a of sensorimototor (M1/S1) cortex near the targeted DMS. To this end, only D1-Cre sections exhibiting a robust intra-striatal loss of Tom^+^ cells were included in the analysis. Due to the high degree of cell overlap in L6, individual optical slices were selected from the middle of each z-stack and analyzed. Annotations were manually drawn to contain 400-450 µm of tissue dorsoventrally in reference to the edge of corpus callosum, stretching mediolaterally from the dorsolateral striatum to the center of the lateral ventricle. Due to lower Tom^+^ intensity in the cortex, Tom^+^ cells were thresholded at 150 and detected using a slightly modified set of criteria [Min. area=60; Max. area=300; Intensity threshold=75]. No cells were discarded due to low fluorescence as long as they met the above inclusion criteria. Annotations were transferred to the contralateral hemisphere and mirror-reflected as before, ensuring measurements were made in near-identical detection conditions.

### Analysis of electrophysiology data

To identify miniature postsynaptic currents, sweeps were analyzed using the open-source program Clampsuite (https://github.com/LarsHenrikNelson/ClampSuite). Sweeps were processed using a zero-phase finite impulse response filter (order = 201, window = 150 samples) with a low pass cutoff at 450 Hz. To automatically identify events, filtered sweeps were deconvolved from a predefined mEPSC or mIPSC template using Fast Fourier Transform and inverted. Specifically, mIPSCs had to satisfy 1) peak amplitude > 6 pA, 2) rise time 0.5-4 ms, and 3) decay time > 4 ms, whereas mEPSCs had to satisfy 1) peak amplitude > 7 pA, 2) rise time 0.5-4 ms, and 3) decay time > 4 ms. Decay time was defined as time to tau, given by peak amplitude divided by 1/*e*. Every identified event was inspected manually. In some cases, mIPSC events were missed by the program and were manually included if they met the above criteria.

**Figure S1.**
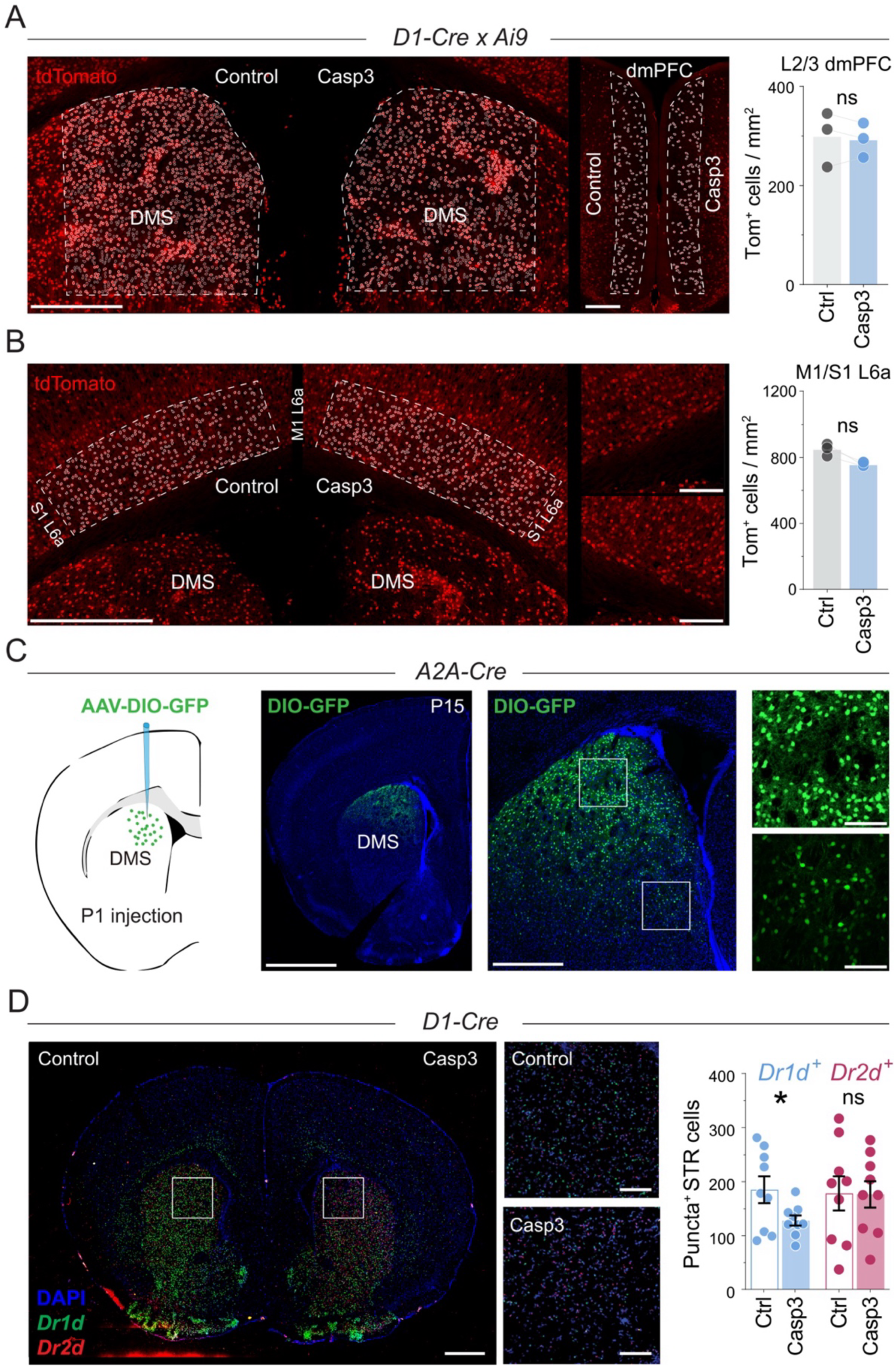
Caspase3 ablation is restricted to DMS and selectively reduces Cre-expressing SPNs. **(A)** Representative maximum projection images from D1-Cre x Ai9^fl/wt^ mice showing tdTomato^+^ cells detected in the DMS of control (left) and Casp3-injected (middle) hemispheres at P14– P18. White outline represents high-intensity and grey outline lower-intensity tdTomato^+^ cells that met detection criteria. Scale bar 0.5 mm. Right: Representative image of tdTomato^+^ cells detected in the superficial (L2/3) dmPFC in D1-Cre x Ai9^fl/wt^ mice injected with Casp3 in the right hemisphere. Scale bar 0.2 mm. Far right: Quantification of tdTomato^+^ cells in L2/3 of the dmPFC in the same mice who received DMS ablation shows no difference between control and Casp3 hemisphere, confirming absence of cortical ablation in the locus of recordings. (Bar plot; Control mean ± SEM = 298.8 ± 32.19 cells/mm^2^, Casp3 mean ± SEM = 292.8 ± 19.87 cells/mm^2^, n = 9 sections from N = 3 mice; Wilcoxon matched-pairs signed rank test, *p* = 0.57; ns = not statistically significant). **(B)** Left: Representative images of tdTomato^+^ cells detected (white outline) in the deep sensorimotor (M1/S1) cortex of D1-Cre x Ai9^fl/wt^ mice who received Casp3 injection in the right DMS. Scale bar 0.6 mm. Right: High-magnification images of control (top) and Casp3-injected (bottom) cortex showing detailed view of L6a superior to the DMS. Scale bar 0.2 mm. Far right: Quantification of tdTomato^+^ cells in L6a of the sensorimotor cortex in mice who received DMS ablation shows a trending albeit nonsignificant decrease in the density of tdTomato^+^ cells in the Casp3 hemispheres, indicating that DMS-targeted ablation did not significantly affect expression of D1-Cre-containg cortical neurons, thus confirming specificity of intra-striatal ablations. (Bar plot; Control mean ± SEM = 849.3 ± 21.52 cells/mm^2^, Casp3 mean ± SEM = 756.6 ± 7.143 cells/mm^2^, n = 7 sections from N = 3 mice; Wilcoxon matched-pairs signed rank test, *p* = 0.25; ns = not statistically significant). **(C)** Left: Schematic of AAV-DIO-GFP injection in the DMS of A2A-Cre mice at P1. Middle and right: Representative slide-scan and maximum projection confocal images of P15 coronal sections showing GFP expression localized to the dorsal striatum with minimal spread to ventral striatal or cortical regions. High-magnification images of boxed regions illustrate restricted AAV spread and absence of GFP^+^ blood vessels, indicating that postnatal injections of Cre-dependent AAVs do not result in recombination in endothelial cells. Scale bars: 1.2 mm, 0.5 mm, and 100 μm, respectively. **(D)** Representative *in situ* hybridization images from D1-Cre mice showing Drd1 (green) and Drd2 (red) mRNA expression in control (left) and Casp3-injected (right) hemispheres. Quantification of labeled DAPI-containing puncta reveals a significant reduction in D1R^+^ but not D2R^+^ cells in the striatum of Casp3-injected animals (Bar plot shows mean ± SEM of puncta-containing DAPI^+^ nuclei in DMS; D1R^+^ Control mean = 184.89 ± 24.64, D1R^+^ Casp3 mean = 128.0 ± 9.6, D2R^+^ Control mean = 178.22 ± 31.43, D2R^+^ Casp3 mean = 176.44 ± 24.6; Wilcoxon matched-pairs signed rank test, *p* = 0.02 for D1R^+^ and *p* = 0.91 for D2R^+^; n = 9 pairs from N = 3 mice). DAPI represents nuclei (blue). Scale bars: 1 mm and 200 μm, respectively.

**Figure S2.**
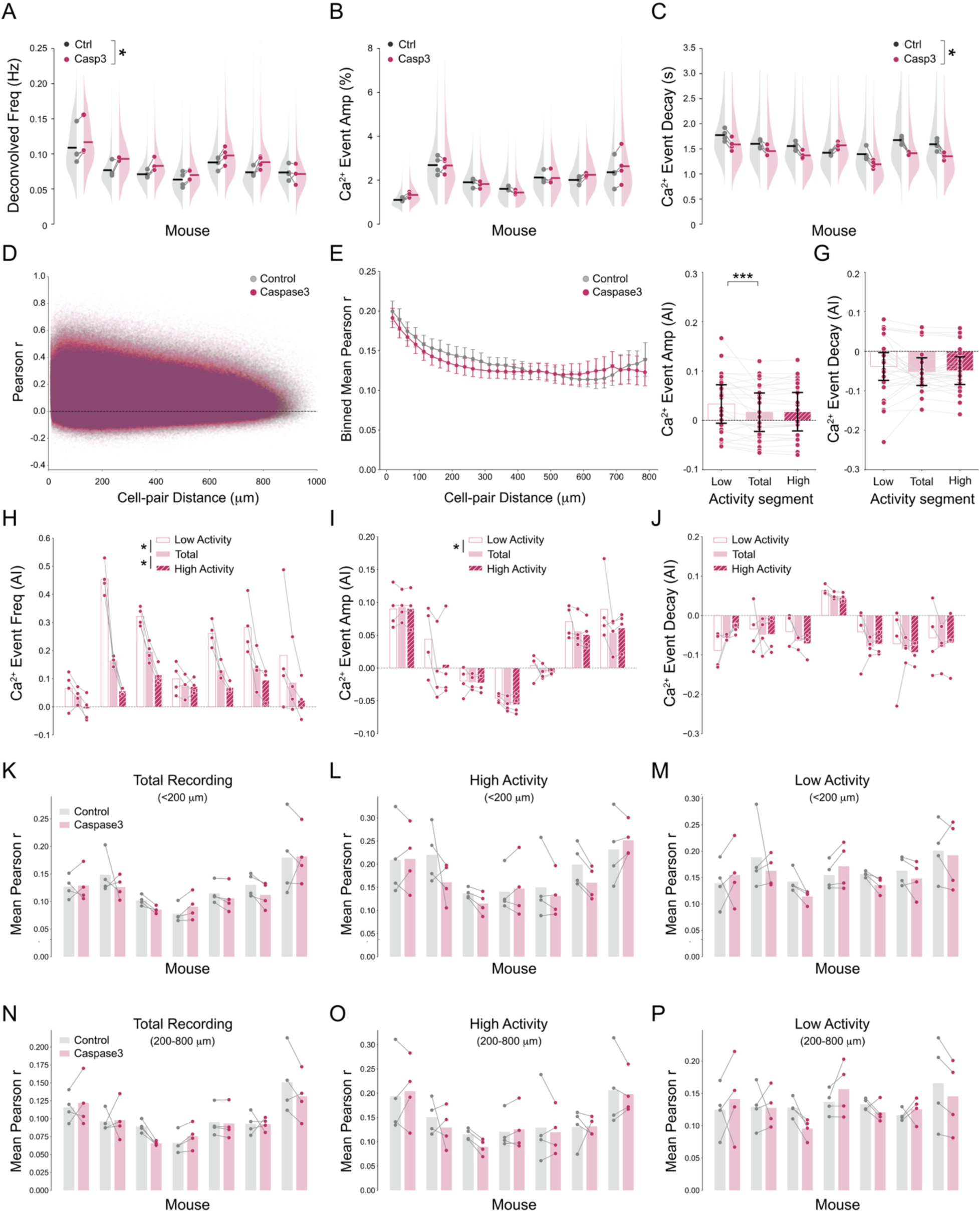
Postnatal ablation of iSPNs increases dmPFC activity. **(A-C)** Analysis of deconvolved impulse frequency of Ca^2+^ events and ΔF/F transient amplitude and decay time per recording in control (gray) versus Casp3ablated (magenta) hemispheres across seven mice. Half-violin plots show each mouse’s bootstrapped distribution of parameters across recordings; dots represent mean of individual recordings; horizontal bars show mouse averages (Deconvolved impulse frequency: Ctrl: 0.079 ± 0.015 Hz, Casp3: 0.088 ± 0.016 Hz, Wilcoxon matched-pairs signed rank test, *p* = 0.031; ΔF/F event amplitude: Ctrl: 1.97 ± 0.51 %, Casp3: 2.03 ± 0.53 %, Wilcoxon matched-pairs signed rank test, *p* = 0.57; ΔF/F event decay time: Ctrl: 1.57 ± 0.13 sec, Casp3: 1.41 ± 0.14 sec, Wilcoxon matched-pairs signed rank test, *p* = 0.046, N = 7 mice). **(D)** Pairwise Pearson correlation coefficients as a function of intercellular distance. Scatter plot represents correlation values by cell-pair distance for all ROIs of all recordings combined. **(E)** Binned average correlation values show pronounced distance dependent changes in pairwise Pearson *r* between short- and long-range neuron pairs. iSPN ablation did not cause changes in pairwise correlation of Ca^2+^ activity. **(F–G)** Asymmetry indices across network activity segments (Low, Total, High) for Ca^2+^ event amplitude and decay time. Bars show estimated marginal means from linear mixed models (LMM; Total as intercept) ± 95 % confidence intervals; magenta dots represent individual recording means; gray lines connect paired hemispheres within each mouse. (Amplitude: Pairwise contrasts: Low vs Total *p* = 9.42 × 10^−6^ [0.01, 0.03]; High vs Total *p* = 0.822 [–0.01, 0.01]. LMM fixed effects: intercept *p* = 0.405 [–0.02, 0.06]; Group variance *p* = 0.123 [–2.69, 22.62]; recording-level variance *p* = 0.0153 [0.48, 4.52]; Decay: Pairwise contrasts: Low vs Total *p* = 0.142 [–0.00, 0.03]; High vs Total *p* = 0.778 [–0.01, 0.02]. LMM fixed effects: intercept *p* = 0.0396 [–0.09, –0.02]; Group variance *p* = 0.171 [–0.66, 3.74]; recording-level variance *p* = 0.0396 [0.05, 1.94]). **(H–J)** Mouse-level asymmetry indices for frequency, amplitude, and decay time across total, high, and low network activity segments. Each group of bars represents a single mouse; bars show mean values for total (solid), high activity (hatched), and low activity (open) segments. Dots represent mean values for individual recordings, and gray lines connect recordings from the same mouse to visualize within-mouse differences across segments. (Frequency: LMM revealed a significant effect of activity segment (LMM, *p* < 0.001). Compared to total, asymmetry indices were significantly increased during low activity (*β* = 0.121, *p* = 4.5 × 10^−8^ [0.079, 0.162]) and significantly decreased during high activity (*β* = –0.057, *p* = 0.007 [–0.099, –0.016]); Amplitude: Asymmetry indices were slightly elevated during low activity segments compared to total (*β* = 0.017, *p* = 0.025 [0.002, 0.031]), but there was no significant difference during high activity (*β* = 0.001, *p* = 0.897 [–0.014, 0.015]); Decay: No significant differences in asymmetry indices were found between activity segments (Low vs Total: *β* = 0.013, *p* = 0.272; High vs Total: *β* = 0.003, *p* = 0.833). **(K–P)** Mouse-level comparison of pairwise signal correlations (Pearson *r*) between control (gray) and Caspase3-ablated (pink) hemispheres across different activity states and spatial scales. Each bar represents the mean correlation between calcium traces for neuron pairs within a given distance range (<200 µm or 200–800 µm), segmented by network activity segment (Total, High, or Low activity). Dots represent individual recordings, and gray lines connect paired control and Caspase3 hemispheres from the same mouse. Short-range correlations (<200 µm): No significant difference in overall correlation between control and Casp3 hemispheres during the full recording (*β* = –0.0071, 95 % CI [–0.0226, 0.0084], *p* = 0.3685). High activity segment showed no significant group difference (*β* = –0.0157, 95 % CI [– 0.0432, 0.0118], *p* = 0.2628). Low activity segment was also not significantly different (*β* = – 0.0094, 95 % CI [–0.0297, 0.0110], *p* = 0.3668); Long-range correlations (200–800 µm): No group effect during the total recording window (*β* = –0.0040, 95 % CI [–0.0156, 0.0077], *p* = 0.5024). High activity segment similarly showed no difference (*β* = –0.0068, 95 % CI [–0.0316, 0.0180], *p* = 0.5916). Low activity segment showed no significant effect (*β* = –0.0031, 95 % CI [– 0.0213, 0.0152], *p* = 0.7433).

**Figure S3.**
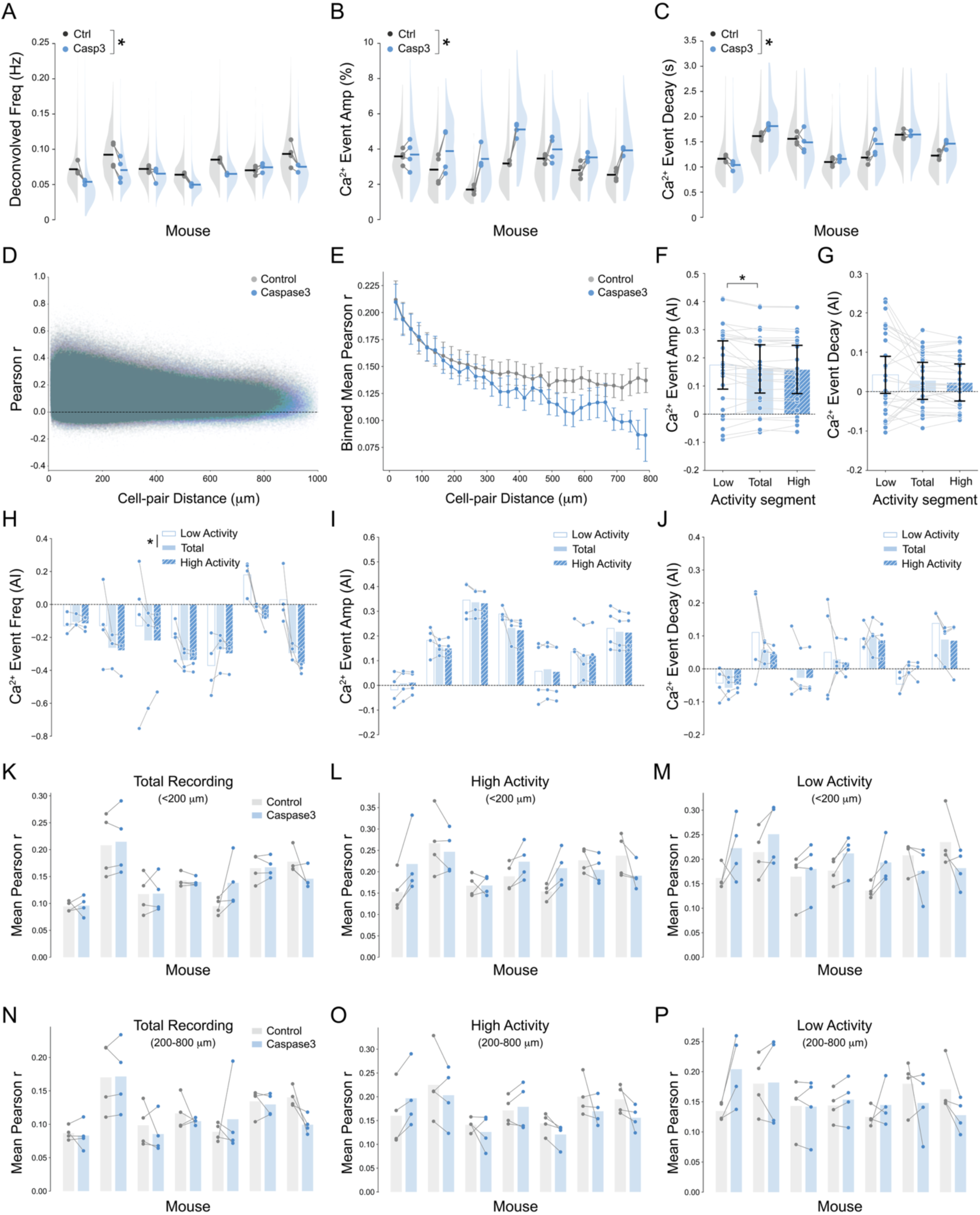
Postnatal ablation of dSPNs reduces dmPFC activity. **(A-C)** Analysis of deconvolved impulse frequency of Ca^2+^ events and ΔF/F transient amplitude and decay time per recording in control (gray) versus Casp3-ablated (blue) hemispheres across seven mice. Half-violin plots represent each mouse’s bootstrapped distribution of parameters across recordings; dots represent mean of individual recordings; horizontal bars show mouse averages (Deconvolved impulse frequency: Ctrl: 0.078 ± 0.012 Hz, Casp3: 0.065 ± 0.010 Hz, Wilcoxon matched-pairs signed rank test, *p* = 0.031; ΔF/F event amplitude: Ctrl: 2.87 ± 0.64 %, Casp3: 3.94 ± 0.56 %, Wilcoxon matched-pairs signed rank test, *p* = 0.016; ΔF/F event decay time: Ctrl: 1.35 ± 0.24 sec, Casp3: 1.44 ± 0.27 sec, Wilcoxon matched-pairs signed rank test, *p* = 0.29, N = 7 mice). **(D)** Pairwise Pearson correlation coefficients as a function of intercellular distance. Scatter plot represents correlation values by cell-pair distance for all ROIs of all recordings combined. **(E)** Binned average correlation values show pronounced distance dependent changes in pairwise Pearson r between short- and long-range neuron pairs. iSPN did not cause changes in pairwise correlation of Ca^2+^ activity. **(F–G)** Asymmetry indices across network activity segments (Low, Total, High) for Ca^2+^ event amplitude and decay time. Bars show estimated marginal means from linear mixed models (LMM; Total as intercept) ± 95 % confidence intervals; magenta dots represent individual recording means; gray lines connect paired hemispheres within each mouse. (Amplitude: Pairwise contrasts: Low vs Total *p* = 0.0197 [0.00, 0.03]; High vs Total *p* = 0.752 [–0.01, 0.01]. LMM fixed effects: intercept *p* = 0.00024 [0.07, 0.25]; Group variance *p* = 0.139 [–7.62, 54.60]; recording-level variance *p* = 0.00761 [2.92, 19.04]; Decay: Pairwise contrasts: Low vs Total *p* = 0.212 [–0.01, 0.04]; High vs Total *p* = 0.730 [–0.03, 0.02]. LMM fixed effects: intercept *p* = 0.255 [–0.02, 0.07]; Group variance *p* = 0.182 [–0.66, 3.46]; recording-level variance *p* = 0.0367 [0.06, 2.04]). **(H–J)** Mouse-level asymmetry indices for frequency, amplitude, and decay time across total, high, and low network activity segments. Each group of bars represents a single mouse; bars show mean values for total (solid), high activity (hatched), and low activity (open) segments. Dots represent mean values for individual recordings, and gray lines connect recordings from the same mouse to visualize within-mouse differences across segments. **Frequency:** LMM revealed a significant effect of activity segment. Compared to total, asymmetry indices were significantly increased during low activity (***β* = 0.106**, *p* = 0.010 [0.025, 0.188]) and not significantly different during high activity (***β* = –0.029**, *p* = 0.488 [–0.110, 0.053]); **Amplitude:** Asymmetry indices did not differ significantly between activity segments (Low vs Total: ***β* = 0.014**, *p* = 0.460 [–0.023, 0.051]; High vs Total: ***β* = –0.002**, *p* = 0.920 [–0.039, 0.035]); **Decay:** No significant differences in asymmetry indices were observed across segments (Low vs Total: ***β* = 0.015**, *p* = 0.356 [–0.017, 0.048]; High vs Total: ***β* = –0.004**, *p* = 0.799 [–0.037, 0.028]). **(K–P)** Mouse-level comparison of pairwise signal correlations (Pearson *r*) between control (gray) and Caspase3-ablated (blue) hemispheres across different activity states and spatial scales. Each bar represents the mean correlation between calcium traces for neuron pairs within a given distance range (<200 µm or 200–800 µm), segmented by network activity segment (Total, High, or Low activity). Dots represent individual recordings, and gray lines connect paired control and Caspase3 hemispheres from the same mouse. **Short-range correlations (<200 µm):** No significant difference in overall correlation between control and Casp3 hemispheres during the full recording (*β* = 0.0027, 95 % CI [–0.0141, 0.0195], *p* = 0.7499). High activity segment showed no significant group difference (*β* = 0.0094, 95 % CI [–0.0148, 0.0336], *p* = 0.4479). Low activity segment was also not significantly different (*β* = 0.0173, 95 % CI [–0.0076, 0.0423], *p* = 0.1737). **Long-range correlations (200–800 µm):** No group effect during the total recording window (*β* = –0.0077, 95 % CI [–0.0233, 0.0080], *p* = 0.3362). High activity segment similarly showed no difference (*β* = –0.0121, 95 % CI [– 0.0349, 0.0106], *p* = 0.2945). Finally, low activity segment showed no significant effect (*β* = 0.0035, 95 % CI [–0.0196, 0.0267], *p* = 0.764).

**Figure S4.**
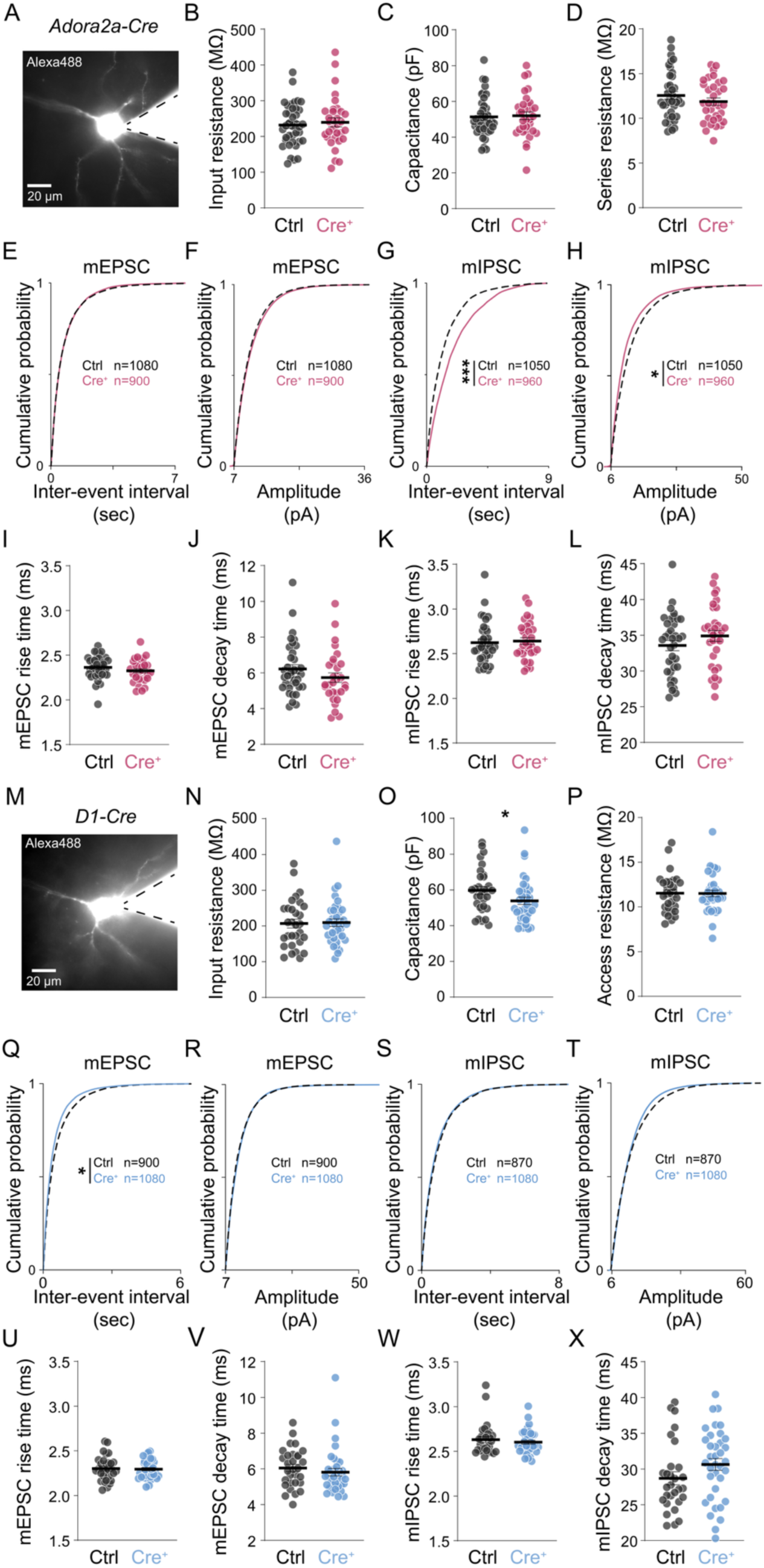
Postnatal ablation of iSPNs disrupts the maturation of GABAergic connectivity in dmPFC. **(A)** Representative image of Alexa-488 filled L2/3 dmPFC pyramidal neuron of A2A-Cre mice in whole-cell configuration. Note the spine-like protuberances along the dendritic arbor and thicker apical dendrite oriented toward the medial surface. Both features were used to include pyramidal neurons in the analysis. **(B)** Input resistance (R_in_) was recorded upon breaking into whole-cell configuration (V_hold_ = −70 mV). Mean R_in_ did not differ between cells sampled from control and A2A-Cre mice (Control = 232 ± 9.37 MΩ, *n* = 39 cells from N = 4 mice; A2A-Cre = 239.6 ± 12.61 MΩ, *n* = 32 cells from N = 3 mice, Mann-Whitney test, *p* = 0.79). **(C)** Mean cell capacitance did not differ between the two groups (Control = 51.28 ± 1.79 pF, *n* = 39 cells from N = 4 mice; A2A-Cre = 51.94 ± 2.19 MΩ, *n* = 32 cells from N = 3 mice, Mann-Whitney test, *p* = 0.69). **(D)** Access resistance (R_s_) was recorded prior to mPSC acquisition, once a cell stabilized ∼4 minutes after breaking into whole-cell configuration (V_hold_ = −70 mV) and monitored continuously to estimate the quality of pipette-cell interface. Only neurons with initial R_s_ < 20 MΩ and final R_s_ < 25 MΩ were analyzed. On average, initial R_s_ did not differ between the two groups (Control = 12.56 ± 0.42 MΩ, *n* = 39 cells from N = 4 mice; A2A-Cre = 11.94 ± 0.41 MΩ, *n* = 32 cells from N = 3 mice; Mann-Whitney test, *p* = 0.35). **(E-H)** Bootstrapped cumulative distribution function (CDF) plots constructed by randomly sampling 30 events from each recorded cell and iterating this resampling process 1000 times with replacement. iSPN ablation did not affect the distribution of inter-event intervals (IEI) of mEPSCs (IEI K-S statistic = 0.01, *p* = 0.99; Control n = 1080 event means from 36 cells, A2A-Cre n = 900 event means from 30 cells) or the distribution of mEPSC amplitudes (Amp K-S statistic = 0.03, *p* = 0.87; Control n = 1080 event means from 36 cells, A2A-Cre = 900 event means from 30 cells). However, iSPN ablation significantly skewed the distribution of mIPSCs toward longer inter-event intervals in the A2A-Cre group (IEI K-S statistic = 0.177, *p* < 0.0001; Control n = 1050 events from 35 cells, A2A-Cre = 960 event means from 32 cells), consistent with a decrease in the mean mIPSC frequency (Fig. 4C). Moreover, the distribution of mIPSC amplitudes was shifted by iSPN ablation to the left, corresponding to a decrease in the amplitude of mIPSCs (Amp K-S statistic = 0.099, *p* = 9.38 × 10^−5^; control n = 1050 events from 35 cells, A2A-Cre = 960 event means from 32 cells) not previously detected at the cell level (Fig. 4G). **(I)** mPSC kinetics were similar across groups: mEPSC rise (Control = 2.364 ± 0.022 ms, n = 36 cells from 4 mice; A2A-Cre = 2.325 ± 0.024 ms; n = 30 cells from N = 3 mice; Mann-Whitney test, *p* = 0.17). **(J)** mEPSC decay (Control = 6.212 ± 0.24, n = 36 cells, A2A-Cre = 5.736 ± 0.27, n = 30 cells, Mann-Whitney test, *p* = 0.18) **(K)** mIPSC rise (Control = 2.623 ± 0.039, n = 36 cells, A2A-Cre = 2.642 ± 0.037, n = 32 cells, Mann-Whitney test, *p* = 0.59). **(L)** mIPSC decay (Control = 33.55 ± 0.71 ms, n = 36 cells, A2A-Cre = 34.91 ± 0.78 ms, n = 32 cells, Mann-Whitney test, *p* = 0.227). **(M)** Representative image of Alexa-488 filled L2/3 dmPFC pyramidal neuron of D1-Cre mice in whole-cell configuration as in (A). **(N)** Input resistance (R_in_) recorded at V_hold_ = −70 mV. Mean R_in_ did not differ between cells sampled from control and D1-Cre mice (Control = 206.8 ± 12.47 MΩ, n = 30 cells from N = 3 mice; D1-Cre = 209.6 ± 10.46 MΩ, n = 32 cells from N = 3 mice, Mann-Whitney test, *p* = 0.89) **(O)** Decreased membrane capacitance in PYRs after dSPN ablation (Control = 59.68 ± 2.33 pF, n = 30 cells; D1-Cre = 53.89 ± 2.07 MΩ, n = 36 cells, Mann-Whitney test, *p* = 0.032) may suggest compensations in somatic and/or neurite volume, possibly suggesting a more complex developmental adaptation not measured in this preparation. **(P)** No difference in access resistance (R_s_) across groups (Control = 11.53 ± 0.396 MΩ, n = 30 cells; D1-Cre = 11.51 ± 0.36MΩ, n = 36 cell; Mann-Whitney test, *p* = 0.9770). **(Q)** CDF analysis revealed a lower mEPSC IEI that is consistent with the elevated mean mEPSC frequency reported in main Figure 4K (IEI K-S statistic = 0.07, *p* = 0.015; Control n = 900 event means from 30 cells, D1-Cre n = 1080 event means from 36 cells), hinting at glutamatergic pre- or post-synaptic adaptations following dSPN ablation. **(R)** Null difference in mEPSC amplitude (Amp K-S statistic = 0.031, *p* = 0.74; Control n = 1080 event means from 36 cells, D1-Cre = 900 event means from 30 cells). **(S)** No differences were observed in the distributions of mIPSC IEIs and **(T)** amplitudes (IEI K-S statistic = 0.028, *p* = 0.84; Amp K-S statistic = 0.041, *p* = 0.34; Control n = 870 events from 29 cells, D1-Cre n = 1080 event means from 36 cells). **(U)** mEPSC rise time did not differ (Control = 2.302 ± 0.025 ms, n = 30 cells; D1-cre = 2.296 ± 0.019 ms, n = 36 cells; Mann-Whitney test, *p* = 0.98). **(V)** mEPSC decay time was preserved (Control = 6.060 ± 0.198, n = 30 cells; D1-Cre = 5.816 ± 0.21, n = 36 cells, Mann-Whitney test, *p* = 0.134). **(W)** mIPSC rise time kinetics were spared (Control = 2.631 ± 0.033, n = 29 cells; D1 = 2.602 ± 0.022, n = 36 cells, Mann-Whitney test, *p* = 0.963). **(X)** mIPSC decay time (Control = 28.71 ± 0.90 ms, n = 29 cells; D1-Cre = 30.65 ± 0.83 ms, n = 36 cells, Mann-Whitney test, *p* = 0.073.)

